# Diacylglycerol kinase-ε is *S-*palmitoylated on cysteine in the cytoplasmic end of its N-terminal transmembrane fragment

**DOI:** 10.1101/2023.06.01.543214

**Authors:** Gabriela Traczyk, Aneta Hromada-Judycka, Anna Świątkowska, Julia Wiśniewska, Anna Ciesielska, Katarzyna Kwiatkowska

**Affiliations:** Laboratory of Molecular Membrane Biology, Nencki Institute of Experimental Biology PAS, 3 Pasteur St., 02-093 Warsaw, Poland

**Keywords:** atypical hemolytic uremic syndrome, cell signaling, diacylglycerol kinase, enzymology/enzyme regulation, kinase activity assay, lipids/chemistry, palmitoylation, phosphoinositides, phospholipids/phosphatidic acid, zDHHC

## Abstract

Diacylglycerol kinase-ε (DGKε) catalyzes phosphorylation of diacylglycerol to phosphatidic acid with a unique specificity toward 1-stearoyl-2-arachidonoyl-*sn*-glycerol which is a backbone of phosphatidylinositol (PI). Owing to this specificity, DGKε is involved in the PI cycle maintaining the cellular level of phosphorylated PI derivatives of signaling activity, and was also found crucial for lipid metabolism. DGKε dysfunction is linked with the development of atypical hemolytic uremic syndrome and possibly other human diseases. Despite the DGKε significance, data on its regulation by co/posttranslational modifications are scarce. Here we report that DGKε is *S-*palmitoylated at Cys38/40 (mouse/human DGKε) located in the cytoplasmic end of its N-terminal putative transmembrane fragment. The *S-*palmitoylation of DGKε was revealed by metabolic labeling of cells with a palmitic acid analogue followed by click chemistry, and with acyl-biotin and acyl-PEG exchange assays. The *S-*acyltransferases zDHHC7 and zDHHC17, and the zDHHC6/16 tandem were found to catalyze DGKε *S-*palmitoylation which also increased the DGKε abundance. Mouse DGKε-Myc ectopically expressed in HEK293 cells localized to the endoplasmic reticulum where zDHHC6/16 reside and in small amounts also to the Golgi apparatus where zDHHC7 and zDHHC17 are present. The Cys38Ala substitution upregulated while hyperpalmitoylation of wild type DGKε reduced the kinase activity, indicating an inhibitory effect of the Cys38 *S*-palmitoylation. Additionally, the substitution of neighboring Pro31 with Ala also diminished the activity of DGKε. Taken together, our data indicate that *S-*palmitoylation can fine-tune DGKε activity in distinct cellular compartments, possibly by affecting the distance between the kinase and its substrate in a membrane.

## INTRODUCTION

Diacylglycerol kinases (DGK) catalyze phosphorylation of diacylglycerol (DAG) to phosphatidic acid (PA). Ten mammalian DGK isoenzymes, α-κ, have been identified and classified into five groups depending on their structure. They all contain two (or three) cysteine-rich conserved homology-1 (C1)-like domains and a catalytic and an accessory domain but their regulatory domains differ (1, 2). DGKε is unique among them - it lacks regulatory domains and is thought to be incorporated into a membrane by a hydrophobic fragment comprising 21 amino acids near its N-terminus (residues 22-42 in human hDGKε, and 20-40 in mouse mDGKε) that is highly conserved in vertebrates (3, 4). DGKε is also distinguished from other DGKs by its specificity toward DAG bearing C16:0 or C18:0 and C20:4 fatty acids at the *sn-*1 and *sn*-2 position, respectively (5–8). The C18:0/20:4 DAG (SAG) is the backbone of phosphatidylinositol (PI) and its phosphorylated derivatives, therefore DGKε is presumed to contribute to the so-called PI cycle together with phosphatidate cytidylyltransferase 2 (CDS2) converting the PA into CDP-DAG (9). This enzymatic cycle serves to rebuild the cellular PI pool following the hydrolysis of PI(4,5)P_2_ in the course of signal transduction by a large number of plasma membrane receptors. Recent data indicate, however, that at a DGKε deficiency, the PI turnover triggered by PI(4,5)P_2_ hydrolysis still occurs, indicating that the specific fatty acyl chain composition of PI and its phosphorylated derivatives can result from an involvement of other DGKs combined with the remodeling of the fatty acyl chains in the so-called Lands’ cycle (10). The interest in DGKε is enhanced by the link detected in mouse models between its activity and the development of Huntington’s disease, seizure susceptibility, and protection from obesity (11–13). Importantly, mutations of *DGKE* are linked with the atypical hemolytic uremic syndrome (aHUS) in humans (14; see https://bibliome.ai/hg19/gene/DGKE for current update). Some of those abnormalities have been correlated with changes in PI(4,5)P_2_ turnover (11, 15).

These features make DGKε of significant interest but the complexity of its regulatory mechanisms is only beginning to be elucidated. Thus, although DGKε contains the C1A and C1B domains resembling those which bind DAG in conventional and novel protein kinase C isoforms, SAG binds to the lipoxygenase (LOX)-like motif of DGKε rather than to those domains (16, 17). On the other hand, the C1 domains can interact with the catalytic domain of DGKε (4, 18) and co-determine its substrate specificity, in analogy to DGKα (8). Our group found recently that the activity and stability of DGKε depend on the zinc finger motif present in the C1B domain. Substitution of any of the cysteines or the histidine of this motif, notably found in some aHUS patients, inactivates the kinase and leads to its proteasomal degradation (19). Finally, the activity and substrate specificity of DGKε are up-regulated allosterically by membrane morphology, i.e., by negatively curved membranes, as found examining purified human DGKε and liposomes of various compositions and sizes (20). The latter suggests that DGKε functioning can depend on its localization in specific regions of cellular membranes.

*S-*palmitoylation is one of the factors that contribute to the accumulation of proteins, both transmembrane and peripheral ones, in selected domains of cellular membranes, including membranes of the endoplasmic reticulum, Golgi apparatus, and the plasma membrane (21–25). This posttranslational modification consists in the attachment of a palmitic acid residue to the sulfhydryl group of cysteine via a thioester bond and is potentially reversible. It is catalyzed by *S-*acyltransferases from the zinc finger DHHC domain containing (zDHHC) family, comprised of 23 enzymes in mammals (26–28). We have recently discovered that DGKε is palmitoylated and the amount of such modified protein increases in Raw264 macrophage-like cells after LPS stimulation (29). It has been established that proteins can be acylated with fatty acids other than palmitic acid, and on rare occasions, the acyl chain can be attached to amino acids other than cysteine (30). Therefore, drawing on recent methodological progress in this work we continued studies on DGKε acylation. The click chemistry-based technique requires metabolic labeling of cells with a fatty acid analogue bearing an alkyne (or azide) moiety at the omega carbon. After cell lysis, the acylated proteins are detected by a “click” reaction between the alkyne (or azide) and an azide (or alkyne) residue of a reporter tag, either biotin or a fluorescent dye. This technique does not discriminate between *S-*acylation of cysteine and the rare *O*- and ε-*N*-acylation at serine or threonine and lysine, respectively (31, 32). On the other hand, the acyl-biotin exchange (ABE) technique relies on the substitution of a thiol-bound fatty acid with a thiol-reactive biotin derivative, hence it only detects *S*-acylation, including *S*-palmitoylation (33). However, it can give false-positive results by detecting proteins forming a thioester linkage with moieties other than fatty acids, like some enzymes of the ubiquitination cascade forming thioester intermediates with ubiquitin (34).

Taking into account the pros and cons of the click-based chemistry and ABE we used both these techniques to study the *S*-palmitoylation of DGKε and identified Cys38/40 (mouse/human DGKε) as the site of this modification. We also found that DGKε can be *S*-palmitoylated by zDHHC6/16, zDHHC7 and zDHHC17, and that this modification down-regulates DGKε activity.

## MATERIALS AND METHODS

### Plasmids

cDNA of mouse DGKε (mDGKε, NM_019505) was obtained and cloned into the pcDNA3.1/Hygro(+) vector (Invitrogen) and fused with a double Myc tag at the C-terminus, as described by Traczyk et al. (19). Plasmid encoding human DGKε (hDGKε, NM_003647) was purchased from OriGene (cat. No. RC219913). The *DGKE* coding sequence was subcloned into the pcDNA3.1/Hygro(+)-2xMyc vector using primers: forward 5’-CTTTCTAGAACCATGGAAGCGGAGAGGCG-3’ and reverse 5’-CTTTCTAGATTCAGTCGCCTTTATATCTTCTTGA-3’ and XbaI restriction enzyme to obtain hGDKε doubly Myc-tagged at the C-terminus. m/hDGKε mutant forms were prepared by site-directed mutagenesis; all wild type and mutant m/hDGKε-Myc constructs were verified by sequencing (Genomed SA). Primers used for *Dgke* and *DGKE* mutagenesis are listed in Supplemental Table S1. A library of pEF-BOS-HA plasmids encoding HA-tagged mouse DHHC1 - DHHC23 *S-*acyltransferases and GST for control were obtained from Prof. Masaki Fukata (National Institute of Physiological Sciences, Okazaki, Japan; 26, 35).

### Cell culture and transfection

HEK293 cells (ATCC CRL-1573™, mycoplasma-free) were plated at 1.2 x 10^6^ in a 6-cm dish (for click, ABE and APE) or 0.5 x 10^6^ in a 3.5-cm dish (for activity measurements and immunofluorescence studies). After 24 h, the cells were transfected with 2 µg (for click and ABE in experiments with no ectopic expression of *S-*acyltransferases) or 850 ng (for activity measurements and immunofluorescence studies) of pcDNA3.1/Hygro(+) plasmids encoding wild type mDGKε or hDGKε or their mutants in 3.3 ml or 1.5 ml DMEM containing 4.5 g/l glucose and 10% FBS, and supplemented with FuGENE HD (Promega) in the ratio 4:1 to the amount of DNA (μl:μg) according to the manufacturer’s instruction. To identify the zDHHC *S*-palmitoylating DGKε, cells (1.2 x 10^6^ in a 6-cm dish) were transfected with 1 µg (ABE) or 2 µg (APE) of pcDNA3.1/Hygro(+) plasmid encoding wild type mDGKε-Myc together with 1 µg (ABE) or 2 µg (APE) of pEF-BOS-HA plasmid bearing one of the 23 *Zdhhc* genes or *Gst* in 3.3 ml DMEM/glucose/FBS using FuGENE HD, as above. In a series of experiments, pcDNA-mDGKε-Myc was co-transfected with pEF-BOS-HA plasmids encoding zDHH6-HA and zDHHC16-HA (1 μg each/dish). Also, when indicated the plasmids encoding mDGKε-Myc or zDHHC-HA were substituted with respective amounts of empty pcDNA3.1 Hygro(+) vector. For the kinase activity measurements in cells co-expressing mDGKε-Myc- and zDHHC17-HA, the proportion of respective plasmids was as for APE, i.e., 850 ng each. Transection was conducted for 24 h when co-expression of mDGKε-Myc with zDHHC-HA was induced (35) or for 48 h for expression of DGKε-Myc alone. In the latter case, the culture medium was exchanged after 24 h for DMEM containing glucose and FBS as above with 0.05 mg/ml streptomycin and 50 U/ml penicillin, and the cell cultivation was continued for additional 24 h. For immunofluorescence studies, after 24 h of transfection, cells were transferred onto coverslips for the remaining 24 h of the cultivation and used for protein localization as described below. In a series of experiments, MG-132 (Merck) was added to the culture medium after 30 h of transfection at 1 μM for the remaining 18 h of the culture. For biochemical assays, the cells were harvested in cold PBS, pelleted and stored at -80°C.

### Click chemistry

After 48 h of transfection with wild type mDGKε-Myc or its mutants, HEK293 cells were incubated with 50 µM 17ODYA (17-octadecynoic acid, Merck) added from stock solution in DMSO or with 0.05% DMSO in DMEM containing 2% charcoal-stripped FBS and 30 mM HEPES for 4 h at 37°C, and processed essentially as described earlier (29). Briefly, harvested and pelleted cells were lysed in 300 µl of lysis buffer containing 0.5% SDS, 0.5% NP-40, 100 mM NaCl, 50 mM phosphate buffer, pH 7.4, 1 mM TCEP (Tris(2-carboxyethyl)phosphine hydrochloride), protease inhibitors (1 mM PMSF, 2 µg/ml aprotinin, 2 µg/ml leupeptin, and 0.7 µg/ml pepstatin), phosphatase inhibitor (1 mM Na_3_VO_4_), protein thioesterase inhibitors (10 µM palmostatin and 0.2 mM HDSF (1-hexadecanesulfonyl fluoride)) and 250 U/ml Benzonase Nuclease (Merck). After 30 min (4°C), the lysates were sheared by passing through a 25-G needle and clarified by centrifugation (4°C, 5 min, 14 500 x *g*), and supernatants were diluted with five volumes of the lysis buffer without the detergents and TCEP, supplemented with 15 µl of Myc-Trap Agarose (with anti-Myc alpaca antibody) (Chromotek, cat. No. yta) and incubated for 3 h at 4°C with end-over-end rotation. Subsequently, samples were washed three times with ice-cold lysis buffer containing 0.05% NP-40 and once with lysis buffer without the detergent. Finally, the agarose beads were suspended in 50 µl of click reaction mixture containing 100 mM NaCl, 10 µM IRDye 800CW-azide (LI-COR, Lincoln), 1 mM TCEP, 100 µM Tris((1-benzyl-4-triazolyl)methyl)amine (TBTA), 1 mM CuSO_4,_ 50 mM phosphate buffer, pH 7.4, and protease inhibitors (cOmplete EDTA-free protease inhibitor cocktail (Roche), 1 mM PMSF, and 0.7 µg/ml pepstatin). In a series of experiments, 5 mM methoxypolyethylene glycol azide (PEG-azide, Merck, cat. No. 689475) was used for the click reaction. After 1 h in the darkness with gentle rotation, samples were washed with lysis buffer containing 0.05% NP-40, and agarose beads were suspended in 40 µl of 2x SDS-sample buffer and heated for 10 min at 95°C. A subset of the 17ODYA-labeled samples were additionally incubated with 1 M hydroxylamine (HXA) for 30 min at 22°C and next diluted twice and incubated for 5 min at 100°C. Proteins were separated by 10% SDS-PAGE and analyzed in an Odyssey CLx Imager (LI-COR) and after that transferred onto nitrocellulose and subjected to immunoblotting.

### Acyl-Biotin Exchange

The acyl-biotin exchange technique was applied to detect *S*-palmitoylation of DGKε essentially as described previously (36), with minor modifications. In brief, HEK293 cells were lysed in 500 μl of 150 mM NaCl, 5 mM EDTA, 1.7% Triton X-100, 4% SDS, cOmplete protease inhibitor cocktail, 2 mM PMSF, 250 U/ml Benzonase Nuclease, 50 mM Tris, pH 7.2, for 15 min at 37°C with shaking, followed by sonication for 60 s (0.3 cycle; amplitude 33%; UP200S Hielscher sonifier). Lysate proteins were precipitated with chloroform:methanol:H_2_O (1:4:3, v:v:v), dissolved in SDS buffer (4% SDS, 5 mM EDTA, 100 mM Hepes, pH 7.4) at 2 mg/ml and incubated with 5 mM TCEP and 20 mM methyl methanethiosulfonate (MMTS) for 15 min at 37°C and 20 min at 50°C, respectively, and precipitated three times as above. For biotinylation, the protein pellet was dissolved in 150 μl of the SDS buffer, diluted 20 times in a buffer containing 0.27 mM HPDP-biotin (ThermoFisher Sci.), 0.27% Triton X-100, and 100 mM Hepes, pH 7.4. Each sample was halved, supplemented either with 1 M hydroxylamine (HXA+) or with 50 mM Tris, pH 7.5 (HXA−) and incubated for 2 h at 20°C with agitation. Proteins were precipitated three times and solubilized as above, 20-μg aliquots were withdrawn as input samples, while 300 μg protein/sample was diluted 20 times in Triton buffer containing 150 mM NaCl, 0.2% Triton X-100, 20 mM Hepes, pH 7.4, 1 μg/ml leupeptin, aprotinin and pepstatin, 1 mM PMSF, 1 mM Na_3_VO_4_, 50 μM phenylarsine oxide, and supplemented with 50 μl of streptavidin-agarose (ThermoFisher Sci.). After 2 h of gentle agitation at 20°C, agarose beads were pelleted and washed 3 times in the Triton buffer without inhibitors. Finally, the beads were suspended in 150 μl of elution buffer (150 mM NaCl, 0.2% SDS, 0.2% Triton X-100, 2% β-mercaptoethanol, 20 mM Hepes, pH 7.5) for 15 min (at 37°C with shaking) and subsequently in 100 μl of 1x SDS-sample buffer for 5 min (at 95°C with shaking). The two eluates were combined, and proteins precipitated with 4 volumes of cold acetone, dissolved in 30 μl of 2 x SDS-sample buffer, denatured for 5 min at 95°C and subjected to 10% SDS-PAGE.

### Acyl-PEG Exchange

To reveal the number of *S*-acylated cysteine residues in DGKε, the palmitic acid residue(s) of mDGKε-Myc were exchanged for polyethylene glycol (PEG) in the acyl-PEG exchange technique (APE, 37) which in principle resembles ABE. For this, HEK293 cells overexpressing mDGKε-Myc together with selected zDHHC-HA or GST for control (24 h) were lysed, the lysates incubated with 5 mM TCEP and 20 mM MMTS with subsequent protein precipitation, as described above. The protein pellet was dissolved in 220 µl of the SDS buffer, protein concentration was adjusted to 2 mg/ml and then diluted 20 times with a buffer containing 0.27% Triton X-100 and 100 mM HEPES, pH 7.4, halved and supplemented or not with 1 M hydroxylamine (HXA+ or HXA−), and incubated for 2 h at 20°C with agitation. Protein was precipitated three times as above and finally dissolved in 100 μl of the SDS buffer. The samples were next supplemented with 100 µl of 4 mM methoxypolyethylene glycol maleimide (PEG-maleimide) (5000; Merck, cat. No. 63187) in the SDS buffer. After 2 h (20°C, with agitation), protein was precipitated 3 times and finally dissolved in 35 µl of the SDS buffer. Protein concentration was determined, samples were supplemented with 12 μl of 4x SDS-sample buffer, incubated for 5 min at 95°C, and subjected in equal amounts to 10% SDS-PAGE.

### DGKε activity determination

The DGKε enzymatic activity was determined using mixed micelle assay according to our previous studies (19). For micelle preparation, 1-NBD-stearoyl-2-arachidonoyl-*sn*-glycerol (NBD-SAG, Cayman Chemical, cat. No. 10011300), SAG (Cayman Chemical, cat. No. 10008650) and 1,2-diacyl-*sn*-glycero-phosphoserine (PS, Merck, cat. No. P7769) dissolved in chloroform:methanol (2:1, v:v, with 30 μg/ml butylated hydroxytoluene as an antioxidant) were mixed at a 5:7 molar ratio of total SAG:PS. The total SAG contained 10% of NBD-SAG. The lipid mixture was dried and resuspended by vortexing (2 min, room temperature) in 4x reaction buffer containing 400 mM NaCl, 80 mM MgCl_2_, 4 mM EGTA, 4 mM DTT, 300 mM octyl-β-glucoside (OG), 200 mM MOPS, pH 7.2, and supplemented with 2 mM ATP. Obtained micelles were used to determine mDGKε-Myc activity in homogenates and detergent lysates of HEK293 cells, and in mDGKε-Myc immunoprecipitates. For homogenization, HEK293 cells 48 h after transfection with mDGKε-Myc alone were suspended in 300 μl of ice-cold homogenization buffer (0.25 M sucrose, 1 mM EDTA, 4 mM EGTA, 1 mM DTT, 20 mM Tris-HCl, pH 7.4, with 1 mM PMSF, 0.7 μg/ml pepstatin, 20 μg/ml aprotinin, and 20 μg/ml leupeptin) and sonicated, as described in (19). For lysis and immunoprecipitation, HEK293 cells 24 h after transfection were used since this time was optimal for co-expression of mDGKε-Myc and zDHHC17-HA, as described above. Cells were lysed essentially as described in (19) at 4°C in 150 µl of lysis buffer supplemented with 1 mM DTT (150 mM NaCl, 20 mM Tris-HCl, pH 7.4, 1% NP-40, 1 mM EDTA, 1 mM DTT, 1 mM PMSF, 20 µg/ml aprotinin and 20 µg/ml leupeptin) and sonicated as above. Then lysates were clarified by centrifugation (4°C, 3 min, 12 500 x *g*) and supernatants were used for the kinase activity measurements or subjected to mDGKε-Myc immunoprecipitation. For the latter, 150 µg of total lysate protein was supplemented with 30 μl of the Myc-Trap Agarose and incubated for 3 h at 4°C with rotation. After washing, the Myc-Trap Agarose with bound mDGKε-Myc was divided in half and one half was resuspended in 40 μl of lysis buffer supplemented with 0.25 M sucrose and 4 mM EGTA and used for the kinase activity assay; the other half was used for immunoblotting (19). For the activity assay, 15 μg of total protein in 40 μl of homogenization buffer or in lysis buffer (containing 0.25 M sucrose and 4 mM EGTA) or half of the mDGKε-Myc immunoprecipitate described above was added to the reaction mixture containing 50 μl of 4 x reaction buffer with micelles and 110 μl of H_2_O. The final concentration of NBD-SAG/SAG in the reaction mixture was 1.45 or 4.46 mol% (0.75 mM or 2.5 mM) and of PS 2.03 or 6.25 mol% (1.05 mM or 3.5 mM). The DGKε activity assay was carried out for 10 min at 24°C with gentle agitation and stopped by the addition of 1 ml of methanol followed by 1 ml of chloroform and 0.7 ml of 1% perchloric acid. After phase separation, lower phase was washed two times with 2 ml of 1% perchloric acid and dried under a nitrogen stream. Lipids were dissolved in chloroform:methanol (2:1, v:v) and separated by TLC on silica gel 60 (Merck) along with a standard, 0.5 -100 pmol of 1-palmitoyl-2-NBD-dodecanoyl-*sn*-glycero-3-phosphate (NBD-PDPA, Avanti, cat. No. 810174), with chloroform:methanol:acetic acid 80% (65:15:5, v:v:v) as mobile phase. NBD-lipids were visualized using G:Box (Syngene) and fluorescence intensity was assessed with ImageJ software using a standard curve drawn for NBD-PDPA and corrected by subtraction of background from samples transfected with empty vector. In parallel, a fraction of the cell homogenates and lysates were supplemented with 2% SDS, vortexed and incubated for 15 min, next supplemented with 1x SDS-sample buffer (final concentration), incubated again for 15 min, heated for 10 min at 95°C and subjected to 10% SDS-PAGE. Also the other half of the mDGKε-Myc immunoprecipitate was suspended in 60 μl of 2 x SDS-sample buffer and heated for 10 min at 95°C to release proteins from the affinity gel for SDS-PAGE. Alongside, 2 - 30 ng of GST-hDGKε (SignalChem, cat. No. D24–10G) was applied onto the gels. After transfer onto nitrocellulose, samples were subjected to immunoblotting with sheep anti-DGKε antibody (R&D Systems) (Supplemental Table S2).

### DGKε localization using immunofluorescence microscopy

HEK293 cells were transfected as described above and after 24 h were transferred onto coverslips (5 × 10^4^ per 15 × 15 mm coverslip) and cultured for additional 24 h. Cells were rinsed twice with ice-cold PBS buffer containing 0.5 mM MgCl_2_ and 1 mM CaCl_2_ and once with APHEM buffer (60 mM PIPES, 25 mM Hepes, 10 mM EGTA, 4 mM MgCl_2_, pH 6.9) and fixed in 4% paraformaldehyde in APHEM for 15 min at room temp. Next, cells were washed with APHEM and incubated with 50 mM NH_4_Cl/APHEM (10 min, room temp.) to block free aldehyde groups. After rinsing with APHEM buffer, cells were permeabilized with 0.05% Triton X-100 in TBS buffer (25 mM Tris-Cl, 130 mM NaCl) for 5 min on ice. To visualize the Golgi apparatus, the protocol described in (38) was used with minor modifications. Cells were permeabilized with 0.005% digitonin in APHEM for 10 min at room temp. and rinsed twice with TBS and incubated with 5% BSA in TBS for 30 min at room temp. Subsequently, cells were incubated overnight at 4°C in 0.2% BSA/TBS with the following antibodies: anti-Myc mouse IgG (Cell Signaling, cat. No. 2276) for mDGKε-Myc staining and anti-STIM1 rabbit IgG (Cell Signaling, cat. No 5668) or anti-golgin-97 rabbit IgG (Cell Signaling, cat. No. 13192) or anti-GM130 rabbit IgG (Cell Signaling, cat. No. 12480). After five washes in 0.2% BSA/TBS, a 1-h incubation with secondary antibodies in 0.2% BSA/TBS was conducted at room temp. The antibodies used were: donkey anti-mouse IgG-Alexa Fluor 647 (ThermoFisher Sci., cat. No. A-31571) and donkey anti-rabbit IgG-FITC (Jackson ImmunoResearch, cat. No. 711-095-152); they were supplemented with 2 µg/ml Hoechst 33342 (Merck, cat. No. B2261). All the antibodies used are listed in Supplemental Table S2. Cells were washed five times with 0.2% BSA/TBS and post-fixed in 2% paraformaldehyde in PBS for 5 min at room temp, washed three times with PBS containing 50 mM NH_4_Cl, once with 0.2% BSA/TBS, once with water, and embedded in Mowiol (Polysciences cat. No. 9002-89-5) containing anti-fading agent DABCO (Merck). Cells were examined under an LSM800 inverted confocal microscope (Zeiss) using a 63 x oil objective (NA 1.4) with scan frame 898×898, speed 8, zoom 1.6, line averaging 2, pixel size 71 nm, and z interval 230 nm. Triple-stained images were obtained by sequential scanning for each channel to eliminate the crosstalk of the chromophores and to ensure a reliable quantification of colocalization. FITC fluorescence was excited at 488 nm, Alexa Fluor 647 at 640 nm, and Hoechst at 405 nm, and detected at 485-540, 637-700, and 400-487 nm, respectively. Images were processed using ImageJ software and finally were exported and maintained as TIFF images for Figure preparation.

For analysis of colocalization about 30 cells were analyzed from two independent transfections. For this, 16-bit z-stacks were used to build 3D structures with Imaris 10.0.1 (Oxford Instruments) software. Next, regions of interest (ROI) were selected on the basis of the staining for DGKε-Myc, STIM1, GM130 or golgin-97 using the surface detection algorithm of Imaris with 220 voxels for GM130 and golgin-97 or 10 voxels for DGKε-Myc and STIM1, and with a manually adjusted threshold. The background threshold values were identical for all images analyzed for the pair of analyzed proteins in a given experiment. To estimate the colocalization of mDGK-Myc in its 3D ROI with a marker protein in the respective ROI, Manders’ colocalization coefficient was used which measures co-occurrence of fluorescence independently of the signal proportionality (39).

### SDS-PAGE and immunoblotting

Proteins were separated by SDS-PAGE and electrotransferred onto nitrocellulose, incubated with antibodies, and processed for detection by chemiluminescence as described previously (19, 29, 40). The following antibodies were used: sheep anti-DGKε (R&D Systems, cat. No. AF7069), mouse anti-Myc (ThermoFisher, cat. No. R950-25), mouse anti-CD71 (transferrin receptor, Santa Cruz Biotechnology, cat. No. sc-32272), rabbit anti-Jak1 (Cell Signaling, cat. No. 3344), mouse anti-actin (BD Transduction Laboratories, cat. No. 612657), rabbit anti-flotillin-2 (Cell Signaling, cat. No. 3436), donkey anti-sheep IgG-HRP (Jackson ImmunoResearch, cat. No. 713-035-003), mouse anti-HA IgG-HRP (Cell Signaling, cat. No. 2999), goat anti-mouse IgG-HRP (Jackson ImmunoResearch, cat. No. 115-035-146) and goat anti-rabbit IgG-HRP (Merck, cat. No. 401315 or Rockland, cat. No. 611-1302). All the antibodies used are listed in Supplemental Table S2. Visualized bands were analyzed densitometrically using the ImageJ program. OD values were corrected by background subtraction, which in the case of click chemistry included samples incubated with DMSO.

## Data analysis

Student’s *t*-test was used for the statistical analysis of data. Data are represented as individual points and mean ± SD.

## RESULTS

### Cys 38/Cys40 is the major site of *S-*palmitoylation of mouse/human DGKε

In order to determine whether DGKε is palmitoylated, we cloned the *Dgke/DGKE* cDNAs and modified them by adding the Myc tag at the C-terminus of the encoded proteins. We decided on such a location of the tag because amino acids 20-40/22-42 (mDGKε/hDGKε) from the N-terminus are predicted to form either a transmembrane or a U-shaped α-helix with Pro31/33 serving as a hinge (9) (Fig. 1A). Mouse DGKε-Myc was overexpressed in HEK293 cells and its palmitoylation was revealed by metabolic labeling with 17ODYA, a palmitic acid analogue subsequently tagged with a fluorescent dye by click-chemistry (Fig. 1B). Strong fluorescence of mDGKε-Myc was observed following its immunoprecipitate separation by SDS-PAGE for 17ODYA-labeled cells, which was virtually absent in a sample from mock-labeled cells (DMSO added instead of 17ODYA) (Fig. 1C, D). An equally negligible mDGKε-Myc fluorescence was observed in another control sample in which the lysate of cells metabolically labeled with ODYA17 was treated with HXA cleaving the thioester bond (Supplemental Fig. S1A-C). The efficient removal of 17ODYA from mDGKε-Myc by HXA indicated that mDGKε undergoes *S*-palmitoylation. In full agreement, an *S-*acylation of mDGKε-Myc was also found with the ABE technique which allows a selective capture of *S-*acylated proteins after substitution of their fatty acid residue(s) with biotin and enrichment on streptavidin beads (Fig. 1E-G). As expected, no mDGKε-Myc was detected in eluates from streptavidin beads incubated with samples not exposed to HXA, which had precluded the biotinylation of originally *S*-acylated proteins (Fig. 1F).

**Figure 1.**
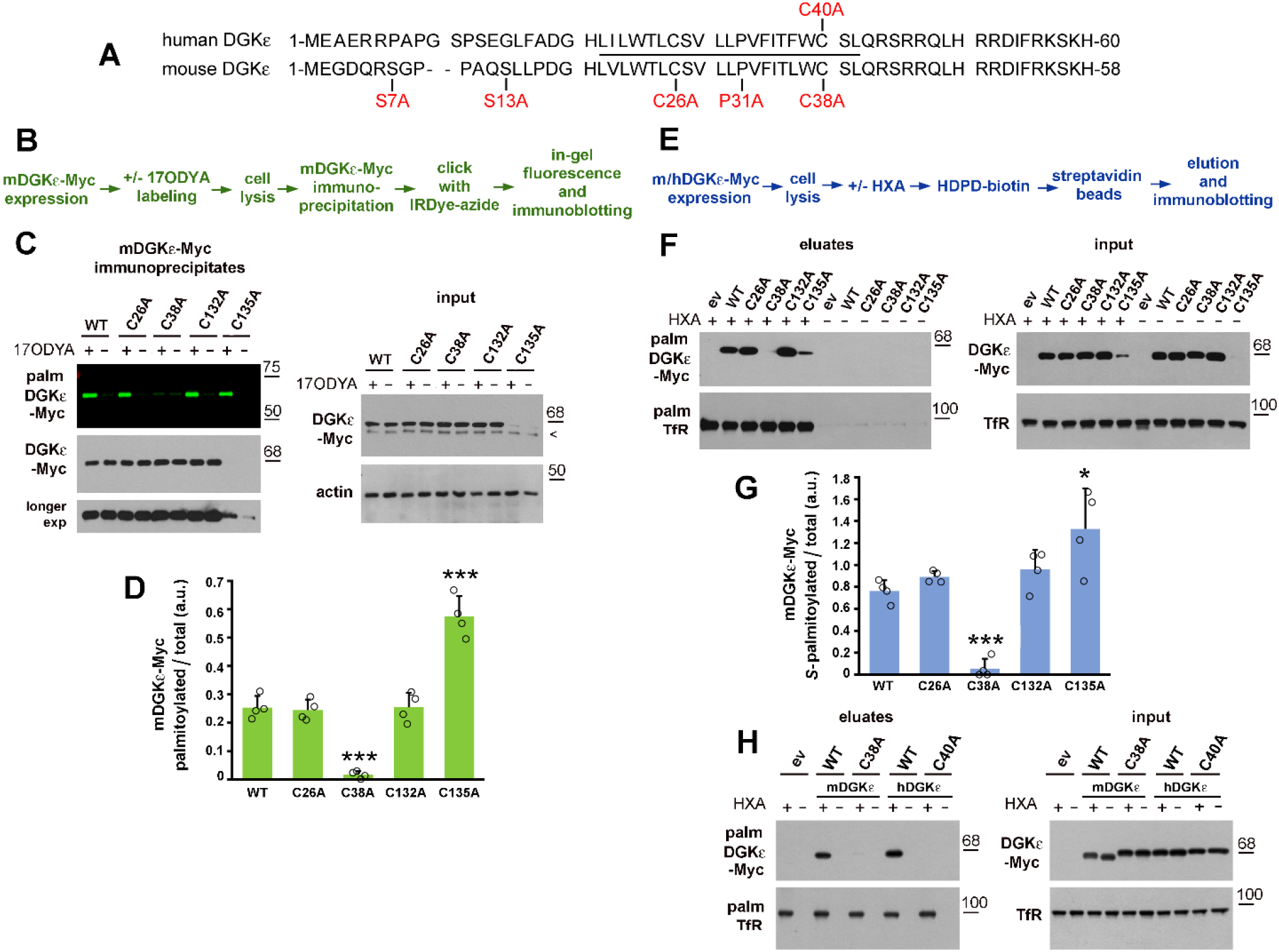
DGKε is *S-*palmitoylated at the cysteine located at the cytosolic end of its N-terminal transmembrane fragment. **(A)** The amino acid sequence of the N-terminal fragment of human and mouse DGKε. Amino acids mutated in this study are indicated by red font. The predicted transmembrane fragment is indicated by the solid line. The complete mDGKε/hDGKε consists of 564/567 amino acids. **(B)** Scheme of the click chemistry procedure. **(C, D, F, G, H)** HEK293 cells were transfected with plasmid encoding wild type mDGKε-Myc **(C, D, F, G)** or hDGKε-Myc (**H**) or their indicated mutant forms. (**C, D)** After 48 h, cells were subjected to metabolic labeling with 50 μM 17ODYA or exposed to 0.05% DMSO carrier as control (−17ODYA) for 4 h and lysed. mDGKε-Myc was immunoprecipitated with anti-Myc alpaca antibody and subjected to click chemistry reaction with IRDye 800CW-azide. **(C, upper panel)** In-gel fluorescence showing mDGKε-Myc labeling with 17ODYA followed by IRDye-azide. **(C, lower panels)** Efficiency of immunoprecipitation determined by immunoblotting with mouse anti-Myc antibody. The content of mDGKε-Myc and actin in input lysates is shown on the right. Arrowhead indicates a band recognized unspecifically by the anti-Myc antibody. **(D)** The extent of mDGKε-Myc palmitoylation. mDGKε-Myc fluorescence was determined by densitometry and normalized against the content of respective Myc-tagged DGKε variant in immunoprecipitates. **(E)** Scheme of the ABE procedure. **(F, G, H)** After 48 h of transfection, cells were lysed and proteins were subjected to the ABE procedure involving treatment with hydroxylamine (HXA+) or not (HXA−), biotinylation, and capture of originally *S-* palmitoylated proteins on strDGKεeptavidin-agarose beads. **(F, H, upper panels)** *S*-palmitoylated mDGKε-Myc **(F)** and hDGKε-Myc **(H)** eluted from the streptavidin-agarose beads revealed with mouse anti-Myc antibody. (**F, H, lower panels)** Transferrin receptor (Tfr), an *S-*acylated protein, eluted from the beads. The content of total mDGKε-Myc, h DGKε-Myc, and TfR in input lysates is shown on the right. **(G)** The extent of mDGKε-Myc *S*-palmitoylation. The content of mDGKε-Myc in eluates and in input lysates was determined by densitometry, normalized against TfR and the ratio of *S*-palmitoylated mDGKε-Myc (eluates) to total mDGKε-Myc (lysates) is shown. WT, wild type; ev, empty vector. Molecular weight markers are shown on the right. Data shown are mean ± SD from four experiments. *, ***, Significantly different at *p =* 0.05 and *p <* 0.001, respectively, from cells expressing wild type DGKε.

To identify the site(s) of the *S*-palmitoylation of DGKε, we focused on its N-terminal region, unique among all DGK isoenzymes and likely forming a transmembrane α-helix (4). We reasoned that cysteine residue(s) located in the membrane-associated fragment are the most likely sites of DGKε *S-*palmitoylation; therefore, Cys26 or Cys38 of mDGKε-Myc were substituted with Ala (Fig. 1A). The Cys38Ala substitution nearly fully abolished the *S*-palmitoylation of mDGKε-Myc, as found with click chemistry and ABE. The trace *S-*acylation represented less than 7% of that found for wild type mDGKε (Fig. 1C, D and F, G). In contrast to Cys38Ala, the Cys26Ala mutation did not affect the *S-*palmitoylation of mDGKε-Myc (Fig. 1C, D and F, G). These data indicate that mDGKε is *S-*palmitoylated nearly exclusively at Cys38. Importantly, also human DGKε was found with the ABE assay to be *S-*palmitoylated and Cys40 (corresponding to Cys38 in mDGKε) was identified as the site of this acylation, as evidenced by its abolishment by the Cys40Ala substitution (Fig. 1H).

Two other cysteine residues, Cys132 and Cys135, located in the C1B domain of mDGKε, had earlier been suggested as sites of its *S*-palmitoylation by a global analysis of the protein *S*-palmitoylation sites in mouse forebrain (41). We therefore mutated either of them and found that the Cys132Ala mutation did not affect the *S-*palmitoylation of mDGKε while Cys135Ala led to its significant augmentation detected with both click chemistry and ABE (Fig. 1C, D, F, G, see also Fig. 2). We have recently found that Cys135 is an element of a zinc finger motif crucial for mDGKε activity and stability and its substitution with Ala leads to proteasomal degradation of the protein (19); indeed, the Cys135Ala mDGKε-Myc variant was present in small quantities which made its detection difficult (Fig. 1C, F, see also Fig. 2). In such conditions the reliability of the quantitation of the ratio of fluorescently- or biotin-labeled to total mDGKε-Myc protein amount is questionable; however, the hyperpalmitoylation of Cys135Ala mDGKε-Myc was confirmed robustly when the amount of the Cys135Ala mDGKε-Myc immunoprecipitate loaded on the gel was increased severalfold to equalize the amounts of Cys135Ala mDGKε-Myc and wild type mDGKε-Myc (Supplemental Fig. S2A-E).

**Figure 2.**
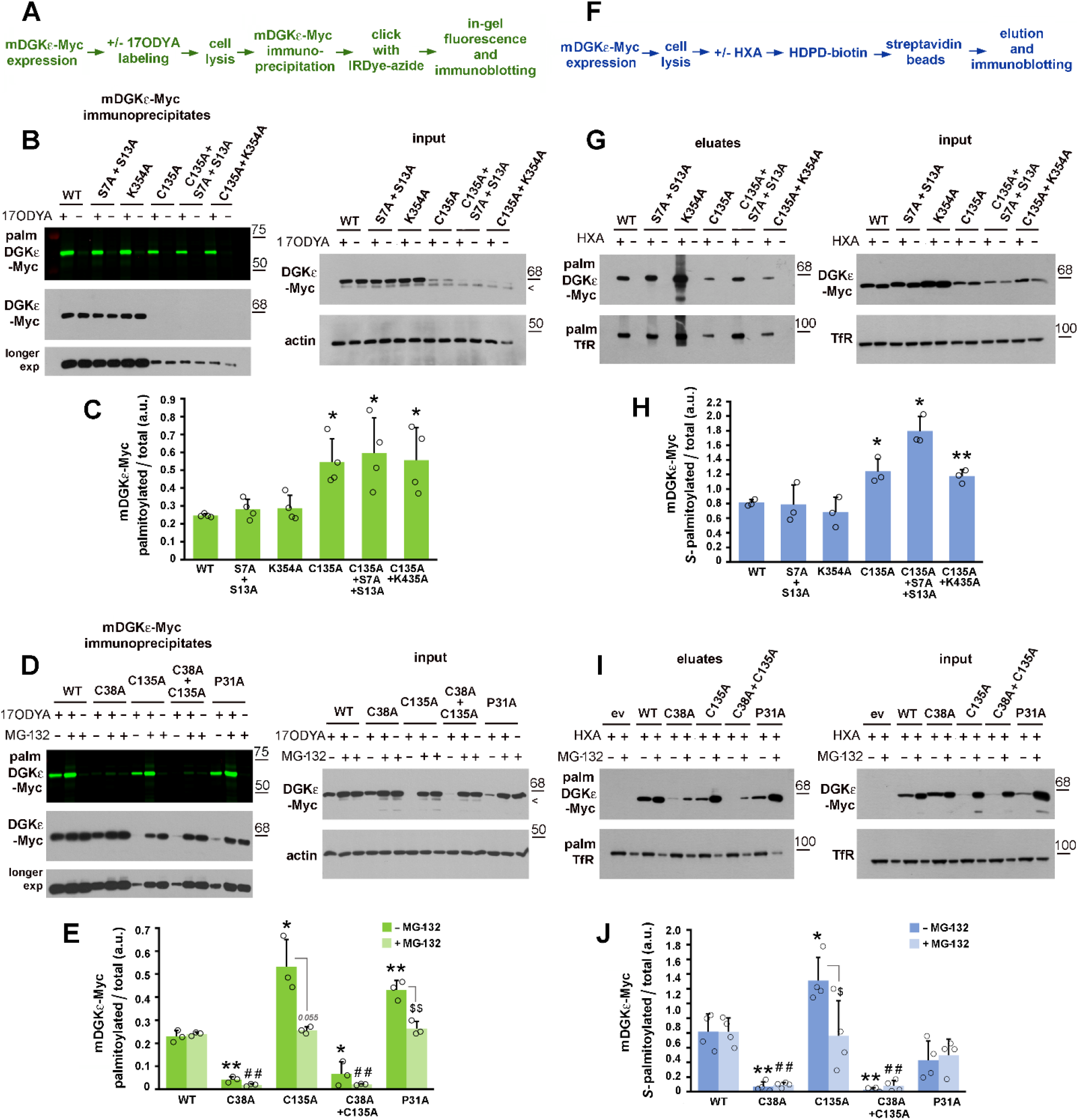
Cys38 is the predominant site of mDGKε *S-*palmitoylation. HEK293 cells were transfected with plasmid encoding wild type mDGKε-Myc or its indicated mutant forms and subjected to click chemistry or ABE. **(A)** Scheme of the click chemistry procedure. **(B-E)** After 48 h, cells were subjected to metabolic labeling with 50 μM 17ODYA or exposed to 0.05% DMSO carrier as control (−17ODYA) for 4 h and lysed. **(D, E)** When indicated, prior to lysis cells were treated with the proteasomal inhibitor MG-132 (1 μM, 18 h). mDGKε-Myc was immunoprecipitated with anti-Myc alpaca antibody and subjected to click chemistry reaction with IRDye 800CW-azide. **(B, D, upper panels)** In-gel fluorescence showing mDGKε-Myc labeling with 17ODYA followed by IRDye-azide. **(B, D, lower panels)** Efficiency of immunoprecipitation determined by immunoblotting with mouse anti-Myc antibody. The content of mDGKε-Myc and actin in input lysates is shown on the right. Arrowheads indicate a band recognized unspecifically by the anti-Myc antibody. **(C, E)** The extent of mDGKε-Myc palmitoylation. mDGKε-Myc fluorescence was determined by densitometry and normalized against the content of respective Myc-tagged DGKε variants in immunoprecipitates. **(F)** Scheme of the ABE procedure. **(G-J)** After 48 h of transfection, cells were lysed and proteins subjected to the ABE procedure involving treatment with hydroxylamine (HXA+) or not (HXA−), biotinylation, and capture of originally *S-*palmitoylated proteins on streptavidin-agarose beads. **(I, J)** When indicated, prior to lysis cells were treated with MG-132 (1 μM, 18 h)**. (G, I, upper panels)** *S*-palmitoylated mDGKε-Myc eluted from streptavidin-agarose beads revealed with mouse anti-Myc antibody. **(G, I, lower panels)** Transferrin receptor (Tfr), an *S-*acylated protein, eluted from the beads. The content of total mDGKε-Myc and TfR in input lysates is shown on the right. In **(I)** MG-132-treated samples were loaded at half the amount of protein from non-treated samples to avoid overloading. **(H, J)** The extent of mDGKε-Myc *S*-palmitoylation. The content of mDGKε-Myc in eluates and in input lysates was determined by densitometry, normalized against TfR and the ratio of *S*-palmitoylated mDGKε-Myc (eluates) to total mDGKε-Myc (lysates) is shown. ev, empty vector. WT, wild type. Molecular weight markers are shown on the right. Data shown are mean ± SD from four (C, J) or three (E, H) experiments. * and ^$^; **, ^##^, and ^$$^; ^###^, significantly different at *p <* 0.05; *p <* 0.01; and *p <* 0.001, respectively, from cells expressing wild type mDGKε-Myc (*, **), wild type mDGKε-Myc in MG-132-treated cells (^##^, ^###^). (^$, $$^) indicate the significance of difference between MG-132-treated and untreated cells.

To confirm further that the mDGKε palmitoylation detected was indeed *S*-palmitoylation, the possibility of *O*-palmitoylation of mDGKε at serine residues Ser7 and Ser13 was investigated. These serine residues could be accessed by acyltransferases of the MBOAT family residing in the endoplasmic reticulum lumen (42, 43) and hypothetically capable of catalyzing *O*-palmitoylation of DGKε in a transmembrane conformation. Both Ser7 and Ser13 were substituted with Ala in wild type or Cys135Ala mDGKε-Myc. These mutations did not affect palmitoylation of wild type mDGKε nor reduced the hyperpalmitoylation of Cys135Ala mDGKε, as found using click chemistry (Fig. 2A-C, Supplemental Fig. S2B, C) and ABE (Fig. 2F-H). These data argue against *O*-acylation of mDGKε, and indirectly confirm that we are dealing with *S*-palmitoylation, in agreement with the susceptibility of the mDGKε – palmitate linkage to HXA treatment.

Next, we substituted Lys354 with Ala, as this residue can be ubiquitinated (9), which could affect mDGKε *S*-palmitoylation, as has been observed for LRP6 (44). Again, no change of mDGKε *S*-palmitoylation was caused by the Lys354Ala substitution, including the hyperpalmitoylation of Cys135Ala /Lys354Ala mDGKε-Myc (Fig. 2B, C and G, H, Supplemental Fig. S2B, C). Notably, the hyperpalmitoylation was nearly abolished in the double Cys38Ala/Cys135Ala mutant, indicating that it resulted from the *S-*palmitoylation of a larger fraction of mDGKε at the “regular” site (Cys38) rather than from acylation of cysteine residue(s) other than Cys38 (Fig. 2D, E and I, J).

Finally, we substituted Pro31 implied to affect the conformation of the N-terminus (4, 9) with Ala to assess its influence on DGKε *S*-palmitoylation. The Pro31Ala substitution decreased the steady-state mDGKε-Myc level (although to a lower extent than did the Cys135Ala substitution) and led to its hyperpalmitoylation; the latter was detected unequivocally with the click chemistry-based approach only (Fig. 2D, E, Supplemental Fig. S2D, E). When the proteasomal activity was inhibited with MG-132, all mDGKε forms were *S-*palmitoylated to a similar extent except for the barely modified Cys38Ala and Cys38Ala/Cys135Ala variants (Fig. 2D, E and I, J). Taken together, the data indicate that Cys38 is the predominant site of mDGKε *S-*palmitoylation, and mDGKε mutated at Cys135 or Pro31 (likely misfolded and rapidly degraded by 26S proteasome) can be hyperpalmitoylated.

### zDHHC7 and zDHHC17 can *S-*palmitoylate DGKε

Having detected the *S-*palmitoylation of mDGKε at Cys38, we aimed to determine which of the 23 mouse zDHHC enzymes could catalyze this reaction. To this end, we overproduced mDGKε-Myc together with each of these enzymes in HEK293 cells and after performing the ABE reaction the amount of mDGKε-Myc eluted from streptavidin beads was compared with that in controls with GST or empty vector in place of zDHHC. The amount of mDGKε-Myc recovered from the beads indicated the extent of its *S-*palmitoylation (Fig. 3A, B, Supplemental Fig. S3A, B). Note that in Fig. 3 the original nomenclature of the *S-* acyltransferases as DHHC is maintained (26, 35), which uses numbering of individual isoenzymes largely coincident with that preferred currently in the zDHHC nomenclature.

**Figure 3.**
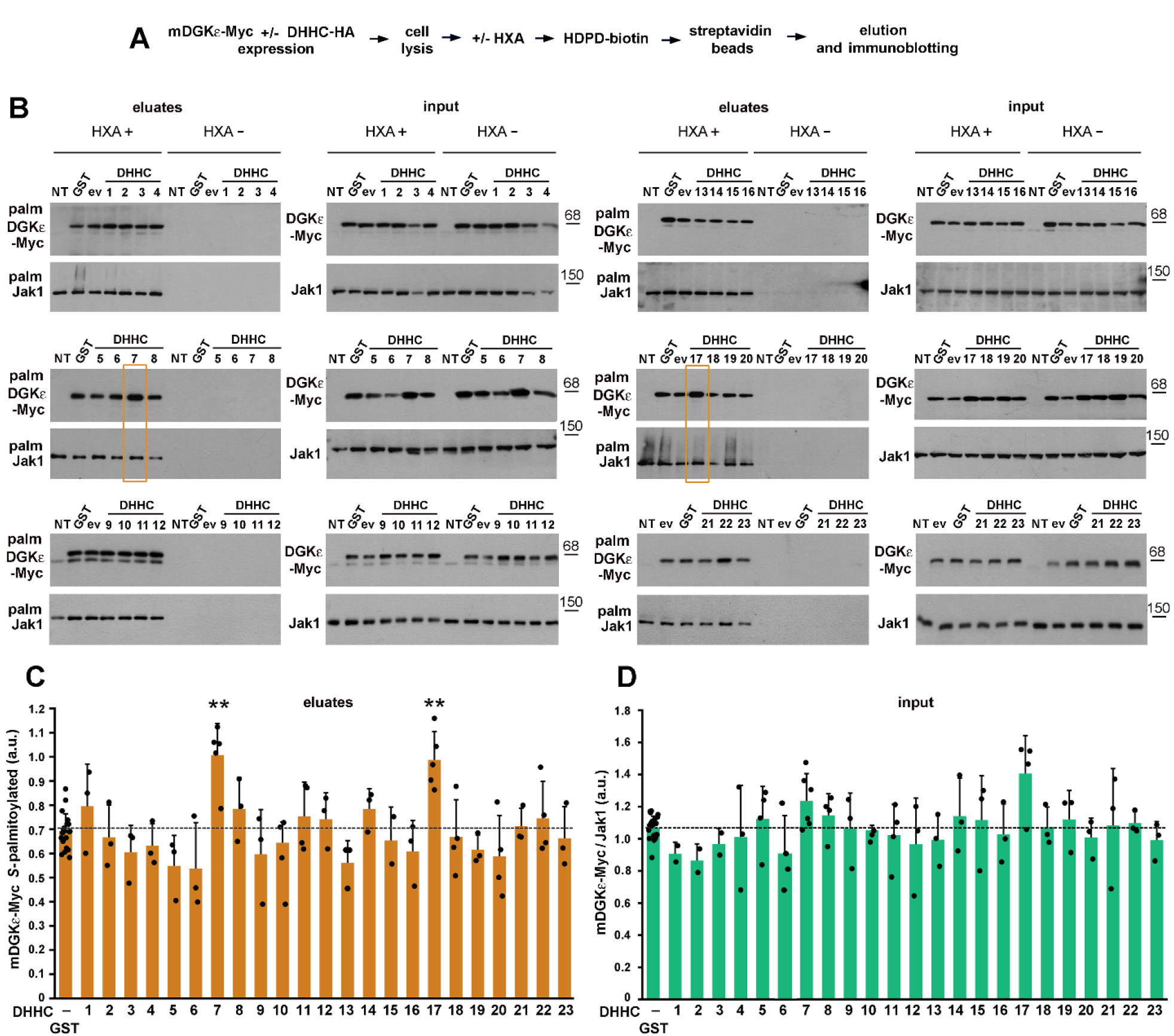
DGKε can be *S*-palmitoylated by zDHHC7 and zDHHC17. HEK293 cells were co-transfected with plasmids encoding wild type mDGKε-Myc and one of the mouse *S-*acyltransferases DHHC1 - 23 tagged with HA, or with GST-HA or empty vector (ev) in controls. After 24 h, cells were lysed and proteins were subjected to the ABE procedure involving treatment with hydroxylamine (HXA+) or not (HXA−), biotinylation, and capture of originally *S-*palmitoylated proteins on streptavidin-agarose beads**. (A)** Scheme of the ABE procedure. **(B)** *S*-palmitoylated mDGKε-Myc (upper panels) and Jak1 (lower panels) eluted from the streptavidin-agarose beads revealed with sheep anti-DGKε and rabbit anti-Jak1 antibody, respectively. The content of mDGKε-Myc and Jak1 in input lysates is shown on the right. NT, cells not transfected. A faint band seen in some panels below mDGKε-Myc represents endogenous hDGKε. Eluates obtained from cells co-expressing mDGKε-Myc and DHHC7 or DHHC17 are boxed in yellow. **(C)** The content of *S*-palmitoylated mDGKε-Myc in eluates was determined by densitometry. **(D)** The content of total mDGKε-Myc in input lysates normalized against total Jak1. Molecular weight markers are shown on the right. Data shown are mean ± SD, experiments in **(D)** were run in duplicates (HXA+ and HXA−). **, significantly different at *p <* 0.01 from cells co-expressing mDGK-Myc and GST-HA. In **(C, D)** the values for GST-expressing cells are averages for all samples tested while the significance of differences was evaluated by comparing values for cells expressing given DHHC with those from GST-expressing cells analyzed on the same gels. DHHC are numbered after Fukata et al. (26, 35), DHHC10, 11, 13, 22 and 23 correspond to zDHHC11, 23, 24, 13 and 25, respectively.

We found that the extent of *S-*palmitoylation of mDGKε-Myc was enhanced significantly in the presence of DHHC7 or DHHC17 (zDHHC7 and zDHHC17, respectively). The amount of mDGKε-Myc eluted from the beads in the presence of either *S-*acyltransferase increased by about 40% in comparison to the GST control (Fig. 3B, C). We found that at the same time also the amount of total mDGKε-Myc protein found in the input cell lysates increased substantially, by 16% in the presence of zDHHC7 and by as much as 31% in the presence of zDHHC17 (Fig. 3B), resembling the effect of zDHHC3, 7, and 17-catalyzed *S*-palmitoylation of sprouty-2 (45).

The results of the analyses are summarized in Fig. 4. We used endogenous Jak1 as a positive control for the recovery of *S-*palmitoylated proteins in all labeled samples (Fig. 3B). Jak1 is *S-*palmitoylated at two cysteine residues (46) and we assured that the level of *S-*palmitoylated endogenous Jak1 eluted from streptavidin beads or the total amount of the protein in input lysates were not affected by the overproduction of zDHHC7 or zDHHC17 (Supplemental Fig. S3C, D). Therefore, the increase of the *S-*palmitoylation of the overproduced mDGK-Myc by zDHHC7 or zDHHC17 was still substantial when normalized to the amount of endogenous *S-*palmitoylated Jak1 found in the eluates (Fig. 4A). Furthermore, we took into account the increase of the total amount of mDGK-Myc (also normalized to Jak1) found in these conditions (Fig. 4B, see also Fig. 3D). As a consequence, the extent of mDGKε-Myc *S-*palmitoylation, i.e., the ratio of the amounts of *S-*palmitoylated mDGKε-Myc and total mDGKε-Myc protein increased marginally in the case of zDHHC7 or remained virtually unchanged for zDHHC17 (Fig. 4C). Taken together, these results show that zDHHC7 and zDHHC17, located in the Golgi apparatus (25), can carry out *S*-palmitoylation of mDGKε (Fig. 4D).

**Figure 4.**
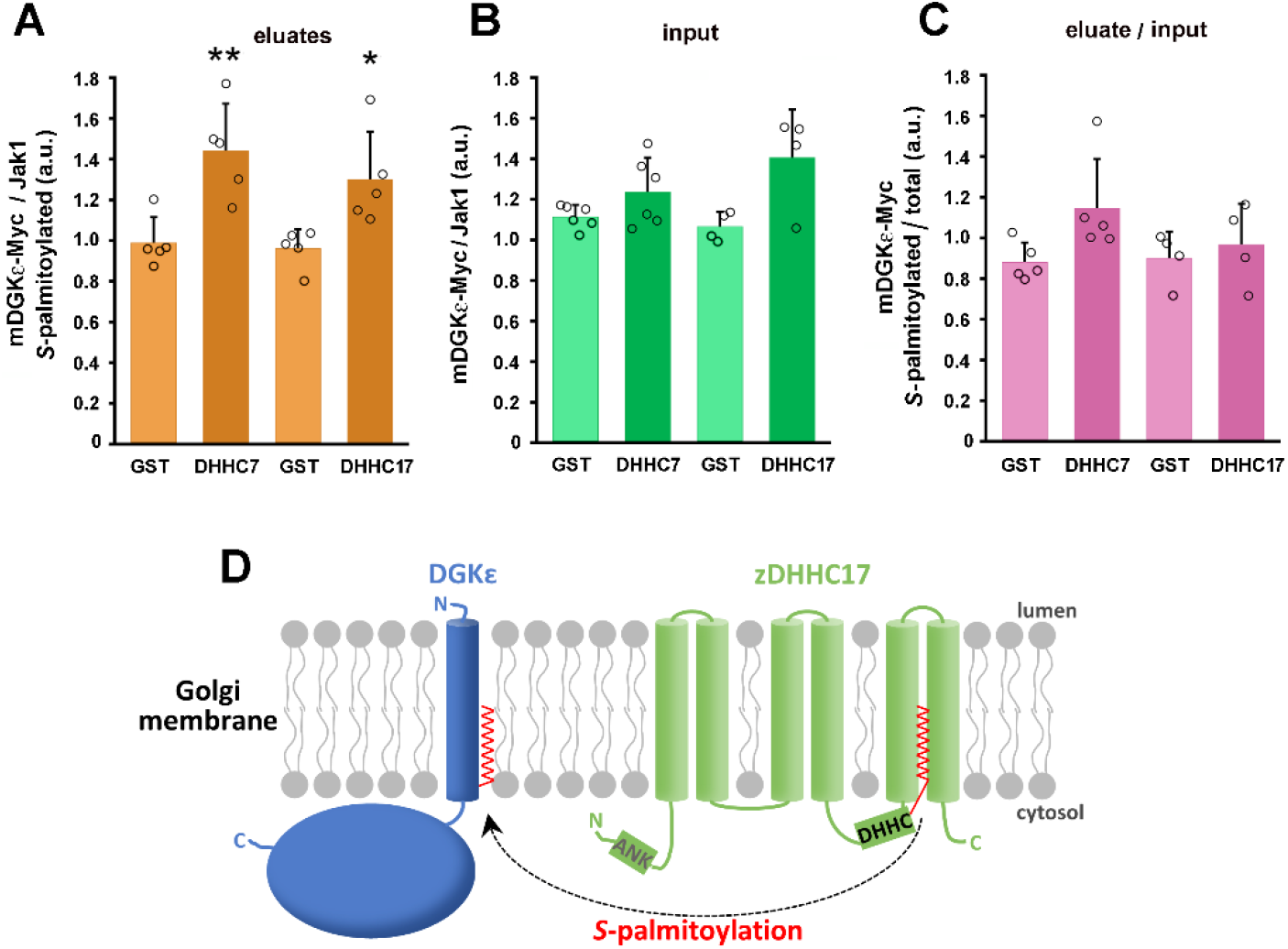
DGKε is *S*-palmitoylated by zDHHC7 and zDHHC17 with a concomitant increase of its abundance. HEK293 cells were co-transfected with plasmids encoding wild type mDGKε-Myc and zDHHC7 or zDHHC17 and analyzed as described in Fig. 3. **(A)** The content of *S*-palmitoylated mDGKε-Myc in eluates was determined by densitometry, as in Fig. 3, and normalized against *S-*palmitoylated Jak1. **(B)** The content of total mDGKε-Myc in input lysates normalized against total Jak1, as in Fig. 3. **(C)** The extent of mDGKε-Myc *S*-palmitoylation in cells co-expressing mDGKε-Myc and zDHHC7 or zDHHC17. The content of mDGKε-Myc in eluates and in input lysates was normalized against Jak1 and the ratio of *S*-palmitoylated mDGKε-Myc (eluates) to total mDGKε-Myc (lysates) is shown. Data shown are mean ± SD from at least four experiments. * and **, significantly different at *p <* 0.05 and *p <* 0.01, respectively, from cells co-expressing mDGK-Myc and GST-HA. **(D)** Scheme of DGKε *S*-palmitoylation by zDHHC17 in the Golgi membrane. *S-*palmitoylation of DGKε by zDHHC7 likely proceeds analogously. All *S-* acyltransferases first bind a palmitic acid residue (red) in the DHHC domain, and then transfer it to a cysteine residue in the substrate protein. zDHHC17 is unique in that it has six transmembrane helices, whereas most zDHHCs have four. It also has an ankyrin-repeat domain (ANK) at the N-terminus involved in interactions with other proteins.

### zDHHC7, 17 and 6/16 *S-*palmitoylate mDGKε at one cysteine residue

Next, we aimed to quantitate the fraction of the *S-*palmitoylated DGKε and verify the number of the *S*-acylation sites. For this, HEK293 cells co-transfected with mDGKε-Myc and selected zDHHCs-HA (or GST for control) were subjected to the APE assay which relies on the substitution of cysteine-bound fatty acyls by the thiol-reactive PEG-maleimide leading to a stable PEGylation of the cysteines (37). The labeling of mDGKε-Myc with PEG slows its gel migration allowing simultaneous detection of the labeled (originally *S-*palmitoylated) and non-modified mDGKε-Myc (Fig. 5A).

**Figure 5.**
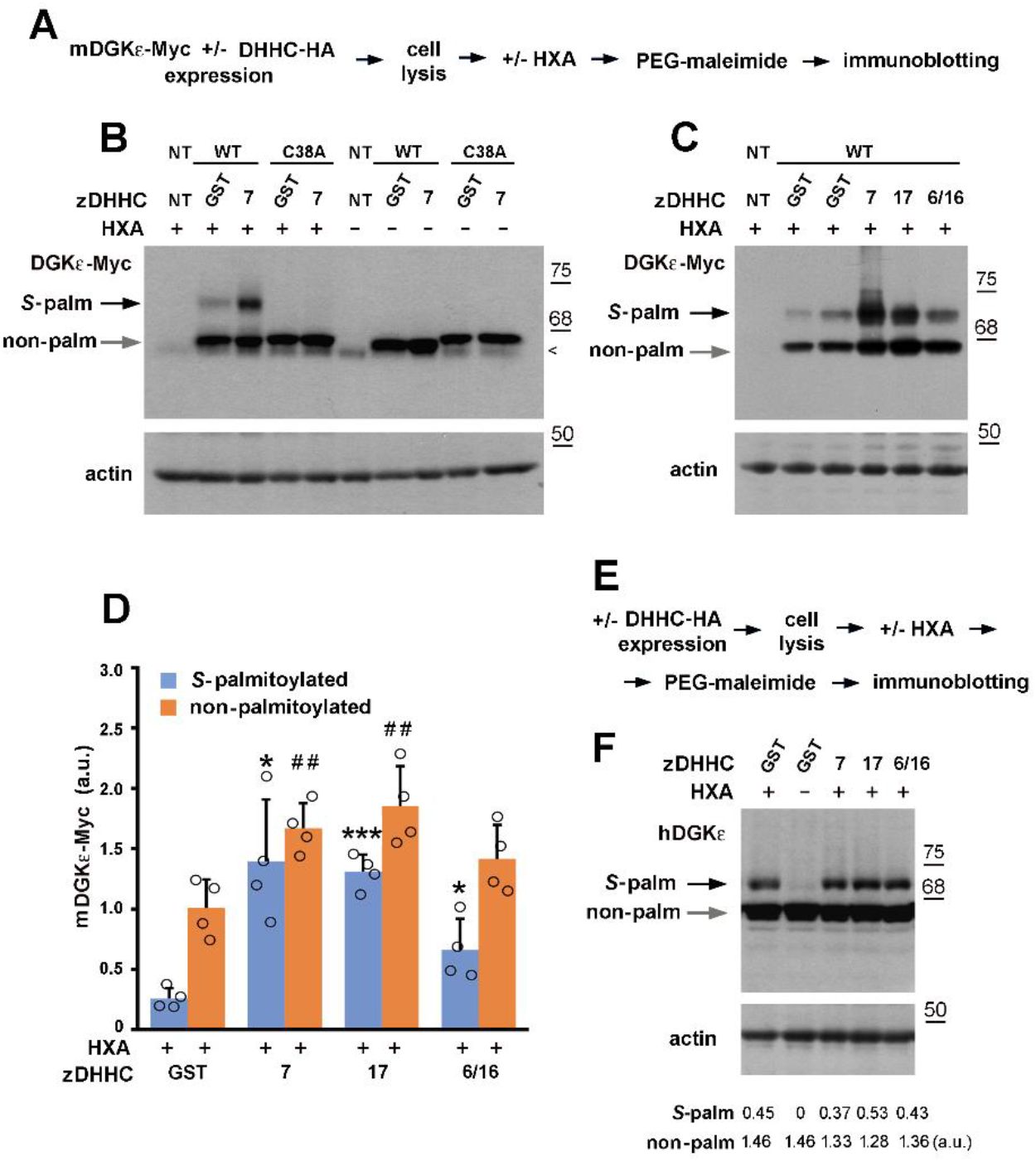
zDHHC7, 17 and 6/16 *S-*palmitoylate DGKε at one cysteine residue. **(A-D)** HEK293 cells were transfected with plasmids encoding wild type mDGKε-Myc or Cys38Ala mDGKε-Myc together with indicated zDHHC-HA or GST-HA in controls. After 24 h, cells were lysed and proteins were subjected to the APE procedure involving treatment with hydroxylamine (HXA+) or not (HXA−) and incubation with PEG-maleimide. **(A)** Scheme of the APE procedure. **(B, C)** mDGKε-Myc (upper panels) and actin (lower panels) in cell lysates revealed with mouse anti-Myc and mouse anti-actin antibody, respectively. NT, cells not transfected. Arrowhead indicates a band recognized unspecifically by the anti-Myc antibody. **(D)** The extent of DGKε *S*-palmitoylation. The content of PEGylated (originally *S*-palmitoylated) and non-palmitoylated mDGKε-Myc was determined by densitometry and normalized against actin. Data are mean ± SD from four experiments. *, ***, significantly different at *p <* 0.05 and *p <* 0.001, respectively, from *S-*palmitoylated form in cells co-expressing mDGKε-Myc and GST-HA; ^##,^ significantly different at *p <* 0.01 from non-palmitoylated form in cells co-expressing mDGKε-Myc and GST-HA. **(E, F)** HEK293 cells were transfected with plasmid encoding indicated zDHHC-HA or GST-HA in controls. After 24 h, cells were lysed and proteins were subjected to the APE procedure. **(E)** Scheme of the APE procedure. **(F)** Endogenous hDGKε (upper panel) and actin (lower panel) in cell lysates revealed with sheep anti-DGKε and mouse anti-actin antibody, respectively. Numbers below blots indicate amount of palmitoylated and non-palmitoylated endogenous hDGKε determined as in **(D)**. WT, wild type. Molecular weight markers are shown on the right.

We focused on zDHHC7 and zDHHC17 but, because DGKε is localized to the endoplasmic reticulum (47, 48), we also included in those experiments zDHHC6 paired with zDHHC16. zDHHC6 is located exclusively in the endoplasmic reticulum and catalyzes *S*-palmitoylation of several local proteins under the control of zDHHC16 (49). To facilitate the subsequent quantitative analyses, in the APE experiments we used twice the amount of the mDGKε-Myc- and zDHHC-encoding plasmids for transfection relative to that used in the earlier ABE quantitative experiments shown in Fig. 3.

We found that in cells expressing mDGKε-Myc with the GST control about 20% of the kinase was modified with PEG reflecting its *S-*palmitoylation by endogenous zDHHC(s) (Fig. 5B-D). Importantly, Cys38Ala mDGKε-Myc was not PEGylated, indicating that Cys38 is the only site of mDGKε *S-* palmitoylation (Fig. 5B). These results were confirmed using a different method - click chemistry between 17ODYA and PEG-azide (Supplemental Fig. S4A-C). In the presence of zDHHC7 or zDHHC17 two phenomena were detected - the amount of *S*-palmitoylated mDGKε-Myc increased by as much as 5-fold bove the GST control, which was accompanied by an increase in the abundance of non-palmitoylated mDGKε-Myc protein by about 65-80% (Fig. 5C, D). As a result, the fraction of PEGylated mDGKε-Myc increased to about 41-46% of the total protein. Bearing in mind the higher levels of ectopically-expressed mDGKε-Myc and zDHHCs in the present experiments, their results are in broad agreement with those of the earlier ones obtained using different methods of analysis (see Fig. 3). In the presence of zDHHC6/16, the increase in the abundance of *S*-palmitoylated mDGKε-Myc was also significant but lower - about 2.5-fold above the GST control. The amount of non-palmitoylated mDGKε-Myc also increased in these conditions (by about 40%), while the fraction of PEGylated mDGKε-Myc accounted for 32% of the total protein (Fig. 5C, D). As expected, when the Cys38Ala mDGKε-Myc was co-expressed with zDHHC7 instead of wild type mDGKε-Myc, no PEGylated (i.e., *S*-palmitoylated) form could be detected (Fig. 5B). In all samples, the mass shift of PEGylated mDGKε-Myc corresponded to the binding of one PEG molecule per mDGKε-Myc molecule, as confirmed by an analysis of flotillin-1 PEGylation (not shown) which is known to be *S-*palmitoylated at a single cysteine (50). However, under conditions favoring strong mDGKε-Myc *S*-palmitoylation, such as its co-expression with zDHH7, an additional faint band appeared above the main one, which could indicate a second site of *S*-palmitoylation (Fig. 5C).

To ensure that the above observations are not unique to the mouse DGKε and are applicable to DGKε of other vertebrates as well, we assessed the *S*-palmitoylation of endogenous hDGKε in HEK293 cells (Fig. 5E, F). Up to 23% of hDGKε was PEGylated in cells not transfected with ectopic zDHHCs, resembling the percentage of modified mDGKε-Myc in corresponding samples. Again, only one site of hDGKε PEGylation was detected (Fig. 5F). The fraction of modified hDGKε did not change in the presence of overexpressed zDHHC7, 17 or 6/16 (Fig. 5F), which suggests that endogenous zDHHC(s) *S*-palmitoylated the kinase to the maximal level.

### *S-*palmitoylation inhibits DGKε activity

The enzymatic activity of wild type, Cys38Ala and Pro31Ala forms of mDGKε-Myc was determined using NBD-SAG-containing mixed micelles in HEK293 homogenates, detergent lysates and mDGKε-Myc immunoprecipitates, in conditions established earlier to ensure a linear dependence of NBD-SAPA production with respect to time and kinase content (19). The NBD-SAPA produced by DGKε was separated by TLC and related to the NBD-PDPA standard (Fig. 6A, D). In parallel, the content of the respective mDGKε-Myc protein in the samples was quantified by immunoblotting using GST-hDGKε as a standard (Fig. 6B, E). The specific activity of wild type mDGKε-Myc in homogenates was about 11 pmol SAPA/min/ng DGKε at 1.45 mol% SAG and did not change when the latter was increased to 4.46 mol% (Fig. 6A-C). The Cys38Ala substitution increased the mDGKε-Myc activity by about 20% at 1.45 mol% SAG and by a further 12% at 4.46 mol% SAG in comparison to the wild type mDGKε-Myc, yet the differences did not the statistical significance (Fig. 6A-C). In contrast, the Pro31Ala substitution reduced the mDGKε-Myc activity significantly by 48% and 42% at 1.45 and 4.46 mol% SAG, respectively (Fig. 6A-C).

**Figure 6.**
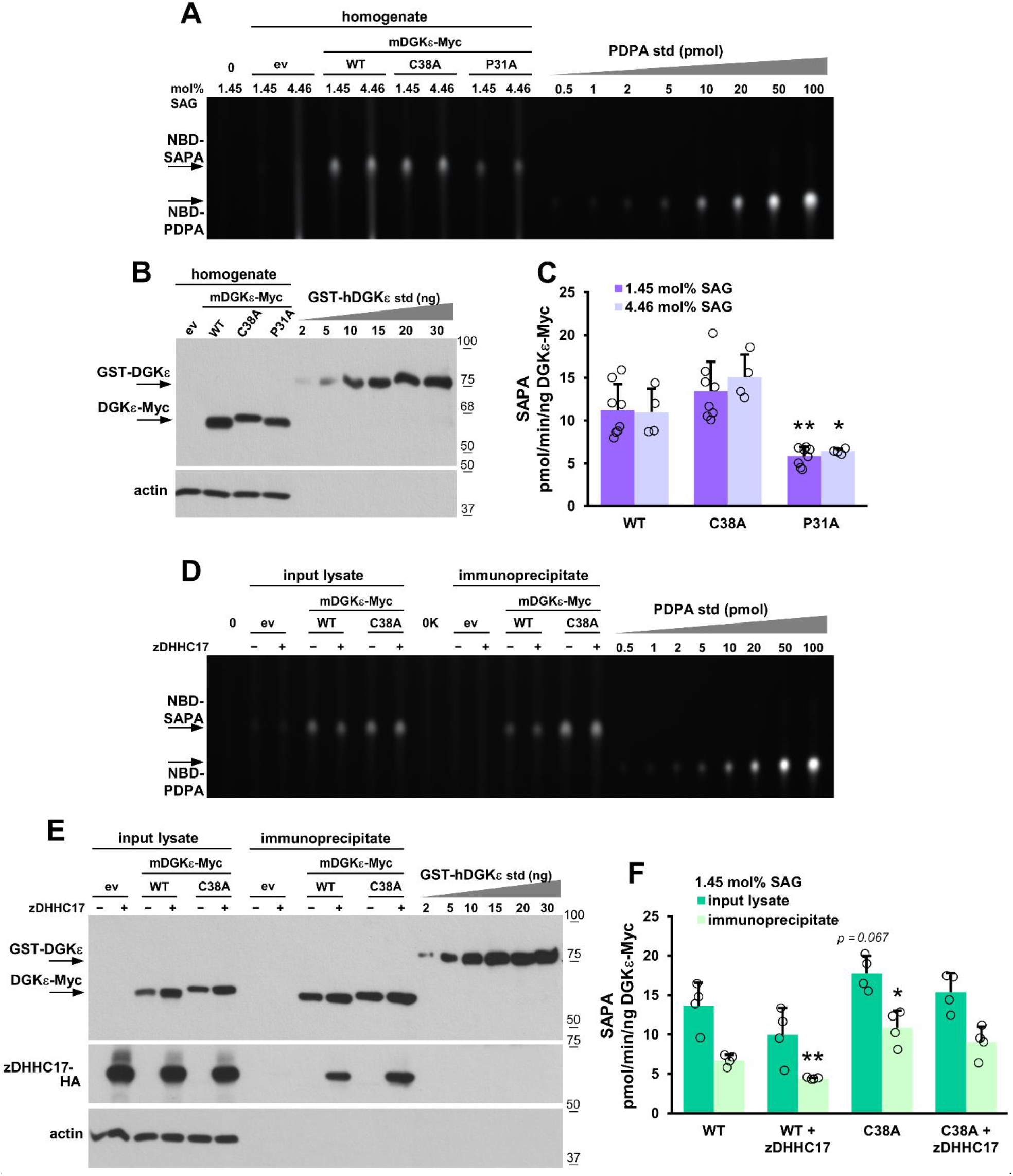
*S*-palmitoylation inhibits DGKε activity. HEK293 cells were transfected with indicated mDGKε-Myc variants or with empty vector (ev) with or without zDHHC17-HA and either homogenized by sonication after 48 h or subjected to cell lysis after 24 h in 1% NP-40 with or without a following mDGKε-Myc immunoprecipitation. The DGKε activity was determined in cell homogenates **(A-C)** or lysates and the mDGKε-Myc immunoprecipitates **(D-F)** using a fluorescence assay with mixed micelles of 1.45:2.03 mol% or 4.46:6.25 mol% NBD-SAG/SAG:PS, as indicated. **(A, D)** Representative TLC separation revealing NBD-SAPA produced in homogenates of mDGKε-Myc-expressing cells (**A**) or in lysates of those cells and in the mDGKε-Myc-immunoprecipitates **(D)**. (0), samples devoid of cell homogenate or lysate, in (0K) supplemented with the Myc-Trap Agarose. Reactions were carried out using 15 μg of total homogenate or lysate protein per sample or using mDGKε-Myc immunoprecipitate obtained from 75 μg of lysate protein. Lipids from 1/50 **(A)** or 1/10 **(D)** of the reaction mixture were separated by TLC. NBD-PDPA is used as a standard, it migrates more slowly on TLC than NBD-SAPA. **(B, E)** Content of indicated overexpressed mDGKε-Myc variants (upper panel) and actin (lower panel) in cell homogenates, lysates and mDGKε-Myc-immunoprecipitates revealed by immunoblotting with sheep anti-DGKε and mouse anti-actin antibody, respectively. GST-hDGKε is used as a standard. In **(E)** also zDHHC17-HA is revealed. Three micrograms of total homogenate protein **(B)** or 10 micrograms of the total lysate protein or 1/3 of the mDGKε-Myc immunoprecipitate **(E)** were applied per lane. WT, wild type. Molecular weight standards are shown on the right. **(C, F)** Specific activity of indicated mDGKε-Myc variants calculated after subtraction of the activity of endogenous DGKs determined in control cells (ev). Data are mean ± SD from eight (1.45 mol% SAG) or four (4.46 mol% SAG) experiments in (C) and from four experiments in (F). * and **, significantly different at *p <* 0.05 and *p <* 0.01, respectively, from samples of wild type mDGKε-Myc analyzed at corresponding mol% SAG.

The results were reproduced when the activity of mDGKε-Myc was measured at 1.45 mol% SAG in HEK293 lysates obtained with 1% NP-40. The activity of Cys38Ala mDGKε-Myc was by about 30% (non-significantly) higher than the activity of wild type mDGKε-Myc (Fig. 6D-E, dark bars in 6F) and that of Pro31Ala mDGKε-Myc activity lower by about 33% (significantly) (Supplemental Fig. S5A-B, dark bar in S5C). Finally, we assessed the activity of the indicted forms of mDGKε-Myc after their immunoprecipitation to minimize a potential influence of unidentified factors present in cell homogenates and lysates. In these conditions, the specific activity of wild type mDGKε-Myc dropped to about 6 pmol SAPA/min/ng DGKε, as found earlier (19). The activity of immunoprecipitated Cys38Ala mDGKε-Myc was significantly higher, by as much as 62% relative to that of wild type mDGKε-Myc, which effect was likely underestimated as only about 20% of the wild type mDGKε-Myc was *S-*palmitoylated (Fig. 6D-E, light bars in 6F). To circumvent this problem, we overproduced wild type mDGKε-Myc together with zDHHC17-HA at conditions inducing its hyperpalmitoylation (see Fig. 5D). Under these conditions, the specific activity of mDGKε-Myc was significantly lower, by 34% in comparison to wild type mDGKε-Myc overproduced without zDHHC17 (Fig. 6D-E, light bars in 6F). Taken together the data indicate that *S-* palmitoylation of DGKε at Cys38 reduces its enzymatic activity. As before, the Pro13Ala mutation reduced the mDGKε-Myc activity measured in its immunoprecipitate (Supplemental Fig. S5A-B, light bars in S5C). Rather unexpectedly, the activity of the nonpalmitoylable Cys38Ala mDGKε-Myc tended to decrease in the presence of zDHHC17-HA (Fig. 6D-E, light bars in 6F). We observed that zDHHC17-HA co-immunoprecipitated with both wild type and Cys38Ala mDGKε-Myc (Fig. 6E). These findings suggest that the interaction of zDHHC17 with mDGKε-Myc alone down-regulates its activity even in the absence of *S-*palmitoylation, while the *S*-palmitoylation of mDGKε-Myc reduces it further.

### mDGKε-Myc is localized in the endoplasmic reticulum and in small amounts in the Golgi apparatus

To analyze the cellular localization of mDGKε-Myc overproduced in HEK293 cells we applied confocal microscopy and markers of the endoplasmic reticulum and the Golgi apparatus. STIM1 was used as a marker of the endoplasmic reticulum membranes, and GM130 and golgin-97 as markers of Golgi membranes (38, 51). To quantitate the distribution of mDGKε-Myc, optical sections of triple-labeled cells (including nucleus labeling) were used to reconstruct 3D images of the mDGKε-Myc-, STIM1-, GM130-or golgin-97-decorated structures (ROI). Next, Manders’ colocalization coefficients were calculated for mDGKε-Myc and each of the marker proteins in the respective ROI. This coefficient scores colocalization regardless of the linear relationship between the intensity of the two signals (39). It allowed us to determine the fraction of a cellular organellum (ROI) decorated by a given marker protein overlapping a DGKε-Myc-positive region (M1) and *vice versa* (M2). As could be expected from earlier studies (47, 48), mDGKε-Myc was abundant in the STIM1-decorated endoplasmic reticulum yielding a white pattern in merged images (Fig. 7A1-C1 and Supplemental Fig. S6A1-C1). The image analysis (Fig. 7D1, Supplemental Fig. S6D1 and Video1) revealed that as much as 80-90% of the STIM1-decorated ROI overlapped with the DGKε-Myc-decorated ROI (Table 1, M1) and 50-60% of DGKε-Myc-positive ROI colocalized with STIM1-positive ROI (Table 1, M2). STIM1 was concentrated in the perinuclear region resembling the pattern observed earlier in resting HEK293 cells (52). DGKε-Myc and STIM1 colocalized to a high extent in this perinuclear part of the endoplasmic reticulum while they tended to be separated toward the cell periphery (Fig. 7C1 and Supplemental Fig. S6C1). This “separate” fraction of DGKε-Myc could be located either in the endoplasmic reticulum devoid of STIM1 or in some other cellular compartment(s).

**Figure 7.**
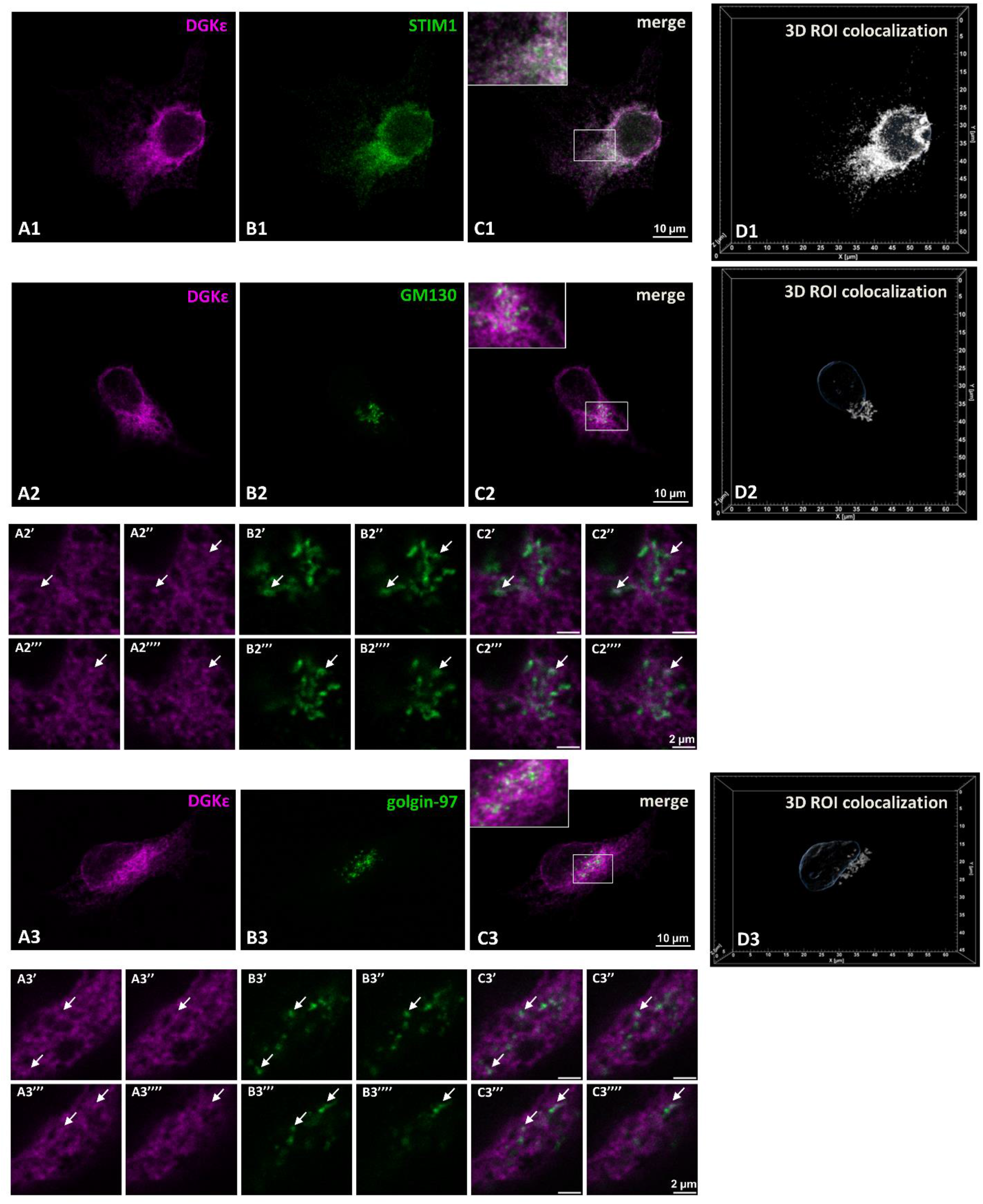
mDGKε-Myc is localized to the endoplasmic reticulum and to the Golgi apparatus. HEK293 cells were transfected with mDGKε-Myc and after 48 h cells were fixed with 4% paraformaldehyde and permeabilized with 0.05% Triton X-100 (for endoplasmic reticulum staining) or with 0.005% digitonin (for Golgi staining). **(A1-A3)** Localization of mDGKε-Myc, **(B1)** STIM1, **(B2)** GM130, **(B3)** golgin-97. **(C1-C3)** Merged images of mDGKε-Myc and the respective marker protein. Colocalized mDGKε-Myc and marker protein appear white. z-Stack images of ten optical sections taken in the middle of a cell are shown. Insets in **(C1-C3)** show enlarged images of marked fragments. **(A2’-A2””)** Four consecutive optical sections through the Golgi showing mDGKε-Myc, **(B2’-B2””)** GM-130, and **(C2’-C2””)** merged image. **(A3’-A3””)** Four consecutive optical sections through the Golgi showing mDGKε-Myc, **(B3’-B3””),** golgin-97, and **(C3’-C3””)** merged images. Arrows point to Golgi fragments with intensive staining of both mDGKε-Myc and the respective marker protein. **(D1-D3)** Reconstructed 3D images of two colocalized ROI positive for mDGKε-Myc and STIM1 **(D1)** or GM130 **(D2)** or golgin-97 **(D3)**. Contours of the nucleus detected by Hoechst 33342 staining are shown in blue. ROI as these were used for quantitative analysis of mDGKε-Myc distribution presented in Table 1. mDGKε-Myc (magenta) was visualized with mouse anti-Myc IgG followed by donkey anti-mouse IgG-Alexa647. STIM1, GM130 and golgin-97 (green) were visualized with rabbit anti-STIM1, anti-GM130, and anti-golgin-97 IgG followed by donkey anti-rabbit IgG-FITC. Additional cells stained according to this protocol are shown in Supplemental Fig. S6.

**Table 1.**
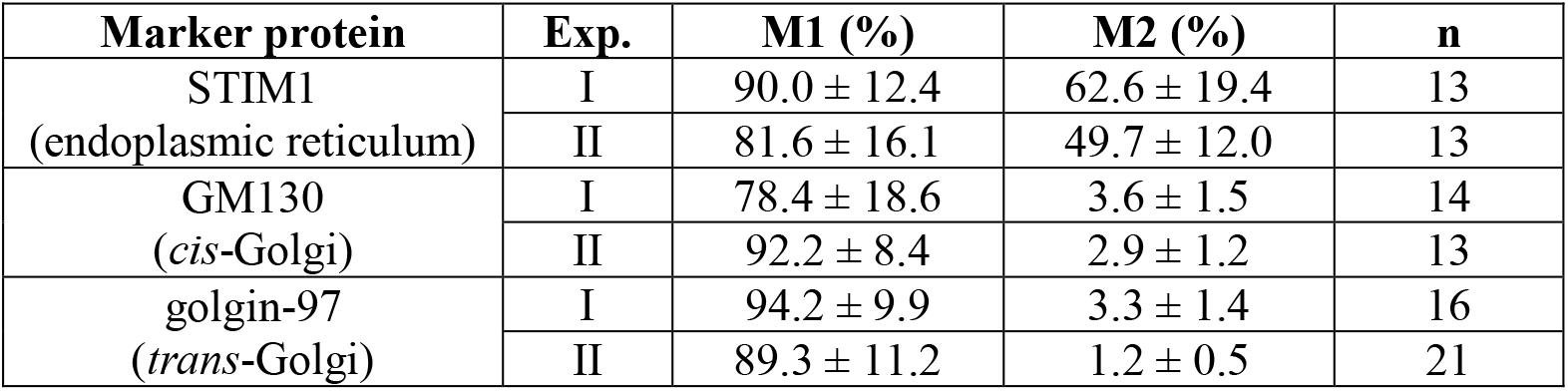
Subcellular distribution of mDGKε-Myc. HEK293 cells were transfected and stained as described in Fig. 7. M1, Percentage of STIM-1, GM130, and golgin-97-positive ROI overlapping mDGKε-Myc-positive ROI. M2, Percentage of mDGKε-Myc-positive ROI overlapping STIM-1, GM130 and golgin-97-positive ROI. Data are mean ± SD from two independent experiments. n = number of cells analyzed.

In order to examine the localization of mDGKε-Myc in the Golgi apparatus, suggested by its zDHHC7- and zDHHC17-catalyzed *S*-palmitoylation, we utilized a protocol optimized for Golgi staining which relies on the cell permeabilization with digitonin at room temperature (38) and used GM130 as a *cis*-Golgi marker (Fig. 7A2-D2, Supplemental Fig. S6A2-D2 and Video 2) and golgin-97 as a *trans*-Golgi marker (Fig. 7A3-D3, Supplemental Fig. S6A3-D3 and Video 3). With this approach, we revealed the presence of some mDGKε-Myc in the Golgi apparatus seen as white spots of its colocalization with GM130 or golgin-97 in Fig. 7C2 and C3 and Supplemental Fig. S6C2 and C3. The white spots can also be observed on merged images of four sequential optical sections shown in Fig. 7C2’-C2”” for mDGKε-Myc and GM130, and in Fig. 7C3’-C3”” for mDGKε-Myc and golgin-97. However, for most regions the intensities of the mDGKε-Myc and GM130 or golgin-97 signals were different from each other not allowing the signals to converge to white. Subsequent image analysis indicated that the fraction of mDGKε-Myc-positive ROI that colocalized with the GM130-positive or golgin-97-positive ROI reached 3-3.5% (Table 1, M2), while as much as 70-90% of either GM130- or golgin-97-decorated Golgi overlapped the mDGKε-Myc-decorated ROI (Table 1, M1). These data indicated that only a small fraction of the total cellular mDGKε-Myc was located in the Golgi apparatus and was distributed through the whole compartment.

We performed a series of control experiments to verify the specificity of the analyzed signals, including the detection of the marker proteins in the absence of the anti-Myc antibody and *vice versa*. No unexpected signal was detected in these conditions (Supplemental Fig. S7 and data not shown) ruling out the possibility of fluorescence bleeding.

## Discussion

The DGK family comprises ten isoenzymes catalyzing phosphorylation of DAG; their activity is likely fine-tuned by co/posttranslational modifications. To date, however, the data on mouse/human DGKε modifications are limited to its ubiquitination at Lys354/357, a scarcity of information surprising in view of the detailed data on the allosteric regulation of DGKε activity found with purified hDGKε (20). Co/posttranslational modifications of all other DGKs are also poorly known, with the exception of their phosphorylation (53). Our earlier study indicated that DGKε can undergo palmitoylation, as revealed by global palmitoylome analysis in Raw264 macrophage-like cells, when also an increase in the amount of palmitoylated mDGKε was detected in cells stimulated with LPS (29). In the present work, we showed that m/hDGKε is *S*-palmitoylated at Cys38/40 and investigated various aspects of this modification in some detail.

Cys38/40 is located at the cytoplasmic side of a putative membrane-bound α-helix of m/hDGKε adopting either a transmembrane topology or a U-shaped intramembrane one with Pro31/33 serving as hinge (9). It has been estimated that in about 95% of single-pass integral membrane proteins their palmitoylation site is located either at a juxtamembrane or an intramembrane cysteine (54), and DGKε fits the first model. Notably, the intramembrane Cys26 of mDGKε was shown here not to be modified. Cys38 is by far the predominant site of DGKε *S-*palmitoylation but perhaps not the only one, as the Cys38Ala substitution eliminated about 93% of the modification detected with click and ABE, but not all of it. The site(s) of this residual *S*-palmitoylation has not been determined nor is it known whether it even occurs in native DGKε. Our preliminary mass spectrometry analyses argue against such a possibility (not shown). There are as many as 28 cysteines in m/hDGKε. So far, no consensus sequence for protein *S*-acylation has been defined conclusively, but Cys72, 73, 75, 87, 88, 149 and 417 of mDGKε (and corresponding Cys residues in hDGKε) are predicted by the SwissPalm database to undergo *S-*palmitoylation with high or medium confidence (https://swissplam.org, 20.10.2023). Notably, all those cysteines except for the latter one are localized to the cysteine-rich domains C1A (amino acids 58/60 - 106/108 in m/hDGKε) and C1B (amino acids 122/125 - 174/177) of DGKε. We have found recently that selected cysteine and histidine residues of the C1B domain of mDGKε form a zinc-finger motif controlling the activity of the kinase and its proteasomal degradation (19). Notably, Cys72/74 and 75/77 belong to an incomplete zinc finger motif lacking a His residue in the C1A domain, while Cys73/75, Cys87/89 and Cys 149/152 are not part of the respective motif in C1A and C1B domain (Fig. 5 in 19), potentially being available for *S-*acylation. A global proteomic analysis of *S*-acylated sites in proteins of the mouse forebrain indicated that Cys132 and Cys135 of the mDGKε C1B domain are modified (39), while our study excluded it. With all the above in mind, we tend to believe that some cysteine residue(s) of the C1A and C1B domains of DGKε can be prone to unspecific labeling at an excess of 17ODYA during metabolic labeling of cells (click technique) and are also not fully reduced/blocked during the ABE procedure, allowing their subsequent biotinylation and false-positive detection.

We identified zDHHC7, zDHHC17 and zDHHC6/16 as candidate mDGKε *S-*acyltransferases using the so-called “Fukata screen” (26, 35) followed by ABE and the APE assay. zDHHC7 and zDHHC17 were more effective than zDHHC6/16 in the model studies. The extent of DGKε *S-*palmitoylation by zDHHC7 or zDHHC17 increased profoundly when the amounts of plasmids encoding these proteins used for transfection were doubled. These results confirm the obvious supposition that the contribution of a particular isoenzyme to the *in vivo* modification of endogenous DGKε would be strongly affected by the possibility of their encounter. The identified zDHHCs differ substantially in their cellular localization, enzymatic activity and also preferences for the acyl-CoA species. zDHHC6 is found exclusively in the endoplasmic reticulum where also DGKε is unequivocally localized, as indicated by the colocalization of mDGKε-Myc with STIM1 in this study and by earlier data obtained for overproduced DGKε as well (47, 48). zDHHC6 catalyzes *S-*palmitoylation of crucial local proteins under the control of zDHHC16 which *S*-palmitoylates zDHHC6 potentially at three cysteine residues of its C-terminal region, with non-palmitoylated zDHHC6 being inactive. The two major *S*-palmitoylated forms of zDHHC6 are, respectively, highly active but short-lived and moderately active but more stable, pointing to a tight regulation of zDHHC6 functioning (49) which was also reflected by a very low level of zDHHC6 overexpression in our studies. On the other hand, zDHHC7 and zDHH17 reside in the Golgi apparatus, as indicated by studies on human and mouse zDHHCs overexpressed in HEK293 and HeLa cells (25, 27). zDHHC7 has a high *S-*acyltransferase activity potentially towards multiple substrate proteins and prefers C14/C16 over C18 acyl-CoA. zDHHC17, with a weaker enzymatic activity, utilizes preferentially the longer C16/C18 rather than C14 acyl-CoA species (55–57). We found that a small fraction of mDGKε-Myc was located in the Golgi apparatus, which could enable its *S*-palmitoylation by zDHHC7 and zDHHC17. We achieved this by using a cell staining protocol preserving the Golgi structure (38) followed by quantitative image analysis. The specificity of the staining procedure and the small amount of mDGKε-Myc found in the Golgi explain the lack of such observations in earlier studies on the cellular location of DGKε. One should bear in mind that the overexpression of zDHHCs and mDGKε could affect their cellular localization, and that 17ODYA used by us for metabolic labeling did not allow discriminating between C16 and C18 acyl chain as natural DGKε *S-*acylation substrates. All the above features are not sufficient to indicate which *S-*acyltransferase isoenzyme is responsible for the DGKε acylation in native conditions. It is of interest in this context that m/hDGKε carries the QP motif found in some zDHHC17 substrates allowing their recognition and binding by the ankyrin-repeat domain of the *S-*acyltransferase (57, 58). zDHHC17 co-immunoprecipitated with mDGKε-Myc in our studies, suggesting, although not proving (59) that DGKε could be a natural target of *S-*acylation by zDHHC17.

The detection of DGKε *S*-palmitoylation, which in HEK293 cells affects about one in four molecules of endogenous hDGKε, raises the question of its physiological significance. Our data indicate that *S*-palmitoylation inhibits the DGKε activity and promotes its accumulation in cells. DGKε has a putative N-terminal transmembrane or re-entrant α-helix and the effects of the *S*-palmitoylation at the cysteine located in the cytoplasmic end of the α-helix can be analogous to those observed for LRP6 or calnexin, both transmembrane proteins of the endoplasmic reticulum *S*-palmitoylated at juxtamembrane cysteines. *S*-palmitoylation of LRP6 (a protein involved in Wnt signaling) is postulated to induce tilting of its transmembrane fragment thereby enabling an adjustment of its effective length to the thickness of the endoplasmic reticulum membrane (60). It also cooperates with monoubiquitination to prevent proteasomal degradation of LRP6 (41). A similar mechanism can explain the observed accumulation of *S*-palmitoylated DGKε and hyperpalmitoylation of Cys135Ala mDGKε-Myc that is avidly degraded by the 26S proteasome (19). The consequences of the *S*-palmitoylation for the conformation of calnexin, an endoplasmic reticulum chaperone which promotes the folding of glycoproteins, are even more complex. The transmembrane fragment of calnexin is also predicted to be tilted in the membrane. Interestingly, the calnexin α-helix is additionally kinked due to the presence of a proline residue somewhat similar to Pro31/33 of m/hDGKε. The *S*-palmitoylation of calnexin by zDHHC6 does not affect the proline-induced kink nor the α-helix tilt in the membrane. However, it is predicted to affect the orientation of its cytosolic fragment with respect to the α-helix axis and can favor the partitioning of calnexin to sheet-like perinuclear structures of the endoplasmic reticulum. As a result, calnexin is properly positioned to capture its client proteins (24). A recent prediction of hDGKε structure indicates that its N-terminal fragment is tilted in the membrane and its conformation determines the distance between the active site and the SAG-bearing membrane. It is proposed that shortening of the distance, facilitating the kinase enzymatic activity, occurs as a result of bending of the transmembrane fragment at Pro33, and such conformational changes can be prevented by the interaction of the positively charged N-terminal amino acids of DGKε with PS in the membrane (4). It seems possible that *S-*palmitoylation of Cys38/40 cooperates with Pro31/33 in a similar manner, preventing the conformational changes of its transmembrane helix and thereby inhibiting DGKε activity. Our data, obtained using immunoprecipitated mDGKε-Myc are consistent with an earlier finding that the whole N-terminal 50-amino-acid-long fragment of hDGKε comprising the transmembrane fragment and the following cluster of positively charged amino acids inhibits the kinase activity, as was determined using an N-terminally truncated purified hDGKε (4). Of note, we found that zDHHC17 co-immunoprecipitated with mDGKε-Myc, including its Cys38A mutant form, and tended to down-regulate (yet insignificantly) the activity of the mutant form of mDGKε-Myc. In this case, the significant inhibition of the wild type mDGKε-Myc activity found in the presence of zDHHC17 can in fact result from a synergy of the zDHHC17 binding DGKε and its *S*-palmitoylation. In addition, the interaction of DGKε with zDHHC17 can increase the cellular abundance of the kinase and this effect was observed in some experiments even for the Cys38Ala mDGKε-Myc mutant form (see Fig. 6).

Our study also shows that the Pro31Ala mutation reduces DGKε activity markedly, in agreement with earlier studies (3). In light of the model discussed above this inhibition may result from hindering the conformational change of the DGKε N-terminus required for its activation. However, in our hands, the Pro31Ala mutation also enhanced the protein degradation in a reproducible manner. We have recently found that certain mDGKε mutants, especially those in its C1B zinc finger, are inactive, likely misfolded, and degraded by the 26S proteasome (19). The Pro31Ala substitution could have a global effect on DGKε folding (hence stability), thereby reducing its enzymatic activity. We base this assumption on the fact that the N-terminally truncated hDGKε had a higher activity but formed insoluble precipitates unless tagged at the N-terminus with His-SUMO (4), indicating an involvement of this DGKε fragment in maintaining proper protein conformation.

Our present study suggests that the *S-*palmitoylation of DGKε can affect its cellular transport and localization. Owing to its strict specificity toward SAG, DGKε has been proposed to contribute to the PI cycle which serves to rebuild PI(4,5)P_2_ level after its hydrolysis triggered by a number of plasma membrane receptors (9, 17). The cycle operates at the plasma membrane-endoplasmic reticulum contact sites formed by STIM1-Orai1 proteins during the receptor activation and requires an involvement of proteins transporting lipids between these two membranes. A discovery of extended synaptotagmins (E-Syts) which mediate bidirectional transport of glycerophospholipids and DAG suggests that the PA of the PI cycle can be formed by DGKε in the endoplasmic reticulum (61). On the other hand, the Nir2 protein transports PA from the plasma membrane toward the endoplasmic reticulum in exchange for PI traveling in the opposite direction thus placing the SAG to PA phosphorylation in the plasma membrane (62, 63). Taking into account that DGKε can be *S-*palmitoylated by zDHHC7 and zDHHC17, one can speculate that it undergoes this modification during its anterograde vesicular trafficking toward the plasma membrane to be kept inactive until its depalmitoylation. *S*-palmitoylation at the cytoplasmic end of transmembrane fragments has recently been identified as a signal guiding diverse transmembrane proteins to the cisternal rims of the Golgi due to its impact on the shape of the transmembrane fragment (its conversion toward a cone-like one). Thanks to such sorting, the acylated proteins are efficiently transported through the Golgi stacks to the plasma membrane (25), and similar trafficking of DGKε *S-*palmitoylated by zDHHC7 or zDHHC17 can be envisioned. A fraction of overexpressed DGKε was localized in the plasma membrane by earlier observations (3), although in our hands mDGKε-Myc was hardly found in this location (unpublished data).

DGKε contributes to lipid metabolism preventing obesity, an activity attributed to DGKε located in the endoplasmic reticulum (13, 64). Furthermore, there are data that question the possibility that DGKε is the sole kinase capable of producing PA required in the PI cycle. Silencing of DGKε alone did not affect the angiotensin-induced PI re-synthesis, clearly pointing to an involvement of other DGKs. On the other hand, the PI re-synthesis was inhibited strongly by the DGK inhibitor R59022 (10) affecting DGKε, α and θ only (65). It has also been found that siRNA silencing of DGKε, and also of DGKα, δ, η, ζ or θ alone did not diminish the 38:4 PA and PI synthesis in unstimulated HEK293 cells (66). To sum up, those two studies indicate that the activity of DGKs other than DGKε, some selective to distinct DAG species (8), is important for the synthesis of PI and its phosphorylated derivatives both in resting cells and during their activation followed by PI(4,5)P_2_ hydrolysis. We found that a stable down-regulation of DGKε using shRNA in resting Raw264 cells decreased the phosphorylation of 38:4 DAG (SAG) to PA by 40% (19), a down-regulation quite substantial *vis-à-vis* the reported lack of an effect of DGKε silencing on PI synthesis found in HEK293 cells. Taken together, the data indicate that, in the absence of DGKε, other DGKs can substitute for its function in *de novo* synthesis and re-synthesis of the major pool of PA/PI.

On the other hand, our data indicating the localization of a small fraction of DGKε in the Golgi apparatus allow us to speculate about its engagement in the maintenance of the local pool of PI(4)P. It has been shown that PI(4)P can be transported from the Golgi toward the plasma membrane (67) and can also be hydrolyzed by most PLC isoforms (68). The PI(4)P-derived SAG lasts longer than the SAG produced by PI(4,5)P_2_ hydrolysis, the major function of which would then be the generation of the IP_3_-mediated Ca^+2^ signal. Furthermore, the PLCε-dependent hydrolysis of PI(4)P at the Golgi is required for the DAG-mediated activation of nuclear PKD during cardiac hypertrophy (68) placing DGKε as a potential terminator of that signal concomitant with the re-synthesis of the local pool of PA/PI(4)P. The DGKε *S-*palmitoylation in the Golgi can facilitate its local retention and increase its abundance, keeping the kinase ready for activation upon depalmitoylation.

## Limitations

The results presented in this study were obtained using mDGKε-Myc overproduced in HEK293 cells. This approach does not undermine the robustness of the detection of the *S*-palmitoylation of m/hDGKε on Cys38/40 or its negative influence on the DGKε activity, although further studies are required to quantitate the kinetic parameters of purified Cys38Ala mDGKε-Myc. Furthermore, the physiological extent of the *S*-palmitoylation of endogenous DGKε by ZDHHC6/16, 7 and 17 can be properly evaluated only by studies of DGKε and the indicated zDHHCs expressed at their native levels. The overexpression of DGKε can in fact result in an underestimation of its Golgi pool. Studies of native DGKε can verify the intriguing possibility of the DGKε involvement in the SAG circulation in the Golgi apparatus which has not been considered so far. although further studies are required to quantitate the kinetic parameters of purified Cys38Ala mDGKε-Myc.

## Data availability

The data that support findings of this study are available from the corresponding author on request.

## Supplemental data

This article contains supplemental data.

## Founding sources

This work was supported by the National Science Centre, Poland (grant number 2018/29/B/NZ3/00407).

## Acknowledgments

We thank Prof. Masaki Fukata from the National Institute of Physiological Sciences in Okazaki, Japan for the library of vectors bearing DHHC1-23 *S-*acyltransferases. We thank Dr. Jan Fronk (retired), formerly at the Faculty of Biology, University of Warsaw for helpful comments, and Dr. Orest V. Matveichuk from our Laboratory for a preliminary study on the identification of zDHHC modifying DGKε. Confocal imaging was performed at the Laboratory of Imaging Tissue Structure and Function, a core facility at the Nencki Institute of Experimental Biology and a part of the infrastructure of the Polish Euro-BioImaging Node.

## CRediT authorship contribution statement

Gabriela Traczyk: Conceptualization, Investigation, Formal analysis, Writing - Review & Editing. Aneta Hromada-Judycka: Investigation, Formal analysis, Writing - Review & Editing. Anna Świątkowska: Investigation, Formal analysis, Writing - Review & Editing. Julia Wiśniewska: Investigation, Writing - Review & Editing. Anna Ciesielska - Investigation, Formal analysis, Writing - Review & Editing; Katarzyna Kwiatkowska: Conceptualization, Supervision, Project administration, Funding acquisition, Writing - Original draft preparation, Writing - Review & Editing.

## Abbreviations

17ODYA: 17-octadecynoic acid
aHUS: atypical hemolytic uremic syndrome
C1: cysteine-rich conserved homology-1
DAG: diacylglycerol
HXA: hydroxylamine
NBD-SAG: 1-NBD-stearoyl-2-arachidonoyl-*sn*-glycerol
NBD-SAPA: 1-NBD-stearoyl-2-arachidonoyl-*sn*-glycero-3-phosphate
NBD-PDPA: 1-palmitoyl-2-NBD-dodecanoyl-*sn*-glycero-3-phosphate
NP-40: Nonidet P-40
OG: octyl-β-glucoside
PA: phosphatidic acid
PI: phosphatidylinositol
PLC: phospholipase C
PS: 1,2-diacyl-*sn*-glycero-phosphoserine
ROI: region of interest
SAG: 1-stearoyl-2-arachidonoyl-*sn*-glycerol or 18:0/20:4-DAG
TCEP: Tris(2-carboxyethyl)phosphine hydrochloride
TLC: thin layer chromatography
zDHHC: zinc finger DHHC domain containing.

