## Supplemental data for "Diacylglycerol kinase-ε is *S-*palmitoylated on cysteine in the cytoplasmic end of its N-terminal transmembrane fragment"

**Supplemental Table S1. Primers used for *Dgke* and *DGKE* mutations**

| mutated form | Primers |
|---|---|
| mDGKε | *Dgke* |
| Ser7Ala Ser13Ala | forward 5'-GGACCAGCGGGCCGGCCCACCCGCCCAGGCCCTGCTCCCTG-3'<br>reverse 5'-CAGGGAGCAGGGCCTGGGCGGGTGGGCCGGCCCGCTGGTCC-3' |
| Cys26Ala | forward 5'-GGCCACTTGGTCCTATGGACGCTGGCCTCCGTGCTGTTGCCGGTGTTC-3'<br>reverse 5'-GAACACCGGCAACAGCACGGAGGCCAGCGTCCATAGGACCAAGTGGCC-3' |
| Pro31Ala | forward 5'-CTCCGTGCTGTTGGCCGTGTTCATCACCTTATG-3'<br>reverse 5'-CATAAGGTGATGAACACGGCCAACAGCACGGAG-3' |
| Cys38Ala | forward 5'-GCCGGTGTTCATCACCTTATGGGCTAGCCTGCAGCGGTCGCGC-3'<br>reverse 5'-GCGCGACCGCTGCAGGCTAGCCCATAAGGTGATGAACACCGGC-3' |
| Cys132Ala | forward 5'-CCGCGGCAACGTCCCCCTGGCCTCTTACTGTGTATTCTGCAGGCAG-3'<br>reverse 5'-CTGCCTGCAGAATACACAGTAAGAGGCCAGGGGGACGTTGCCGCGG-3' |
| Cys135Ala | forward 5'-CAACGTCCCCCTGTGCTCTTACGCTGTATTCTGCAGGCAGCAGTGTGGC-3'<br>reverse 5'-GCCACACTGCTGCCTGCAGAATACAGCGTAAGAGCACAGGGGGACGTTG-3' |
| Lys354Ala | forward 5'-GTTCAGGTAACAAATGCCGGATACTACAATTTAAG-3'<br>reverse 5'-CTTAAATTGTAGTATCCGGCATTTGTTACCTGAAC-3' |
| hDGKε | *DGKE* |
| Cys40Ala | forward 5'-GCCGGTGTTCATCACCTTCTGGGCTAGCCTCCAGCGGTCGCGC-3'<br>reverse 5'- GCGCGACCGCTGGAGGCTAGCCCAGAAGGTGATGAACACCGGC-3' |

**Supplemental Table S2. List of used antibodies**

| Specificity of antibody | Host | Company | Catalog No. | Dilution | Application |
|---|---|---|---|---|---|
| Primary antibodies | | | | | |
| Actin | Mouse IgG1κ | BD Transduction Lab. | #612657 | 1:10 000-1:15 000 | IB |
| CD71 | Mouse IgG1κ | Santa Cruz Biotechnology | sc-32272 | 1:2000 | IB |
| DGKε | Sheep IgG | R&D Systems | #AF7069 | 1:1000 | IB |
| Flotillin-2 | Rabbit IgG | Cell Signaling Techn. | #3436 | 1:3000-1:8000 | IB |
| GM130 | Rabbit IgG | Cell Signaling Techn. | #12480 | 1:500 | IF |
| golgin-97 | Rabbit IgG | Cell Signaling Techn. | #13192 | 1:200 | IF |
| Jak1 | Rabbit IgG | Cell Signaling Techn. | #3344 | 1:1000 | IB |
| Myc | Mouse IgG1 | ThermoFisher Scientific | #R950-25 | 1:1000-1:4500 | IB |
| Myc | Mouse IgG2a | Cell Signaling Techn. | #2276 | 1:500 | IF |
| STIM1 | Rabbit IgG | Cell Signaling Techn. | #5668 | 1:500 | IF |
| Secondary antibodies | | | | | |
| sheep IgG-HRP | Donkey | Jackson ImmunoResearch | #713-035-003 | 1:10 000 | IB |
| mouse IgG-HRP | Goat | Jackson ImmunoResearch | #115-035-146 | 1:6000-1:30 000 | IB |
| rabbit IgG-HRP | Goat | Merck | #401315 | 1:8000-1:10 000 | IB |
| rabbit IgG-HRP | Goat | Rockland | #611-1302 | 1:8000-1:10 000 | IB |
| HA IgG-HRP | Mouse | Cell Signaling Techn. | #2999 | 1:1000-1:5000 | IB |
| mouse IgG-Alexa Fluor 647 | Donkey | ThermoFisher Scientific | #A-31571 | 1:500 | IF |
| rabbit IgG-FITC | Donkey | Jackson ImmunoResearch | #711-095-152 | 1:300 | IF |
| DNA staining | | | | | |
| Hoechst 33342 | Merck | | #B2261 | 2 µg/ml | IF |

IB, immunoblotting; IF, immunofluorescence

(1) Szymańska E, Sobota A, Czuryło E, Kwiatkowska K. Expression of PI(4,5)$P_2$-binding proteins lowers the PI(4,5)$P_2$ level and inhibits FcγRIIA-mediated cell spreading and phagocytosis. (2008) *Eur. J. Immunol*. 38:260-272. doi: 10.1002/eji.200737

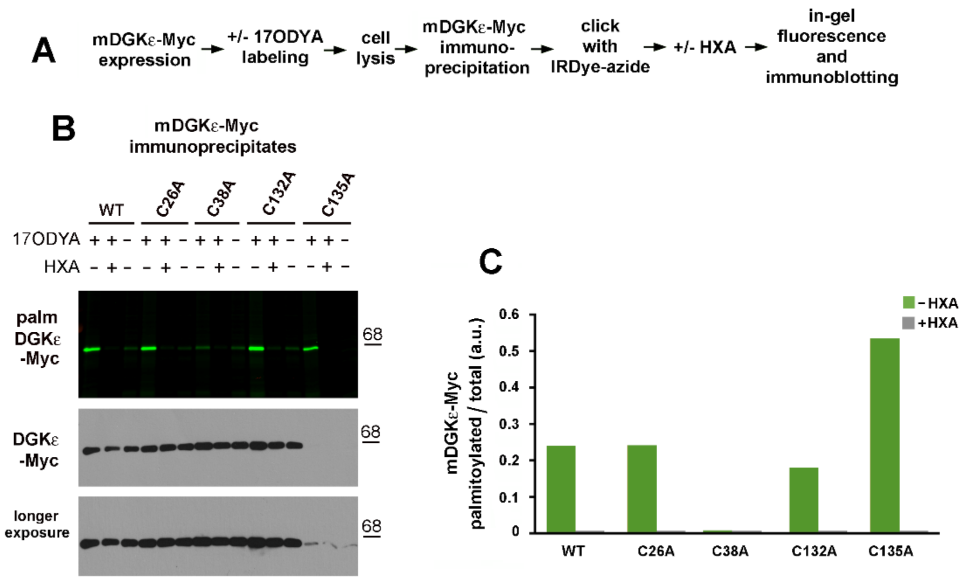


**Supplemental Figure S1. Hydroxylamine removes 17ODYA attached to mDGKε-Myc.** HEK293 cells were transfected with plasmid encoding wild type mDGKε-Myc or its indicated mutant forms. After 48 h, cells were subjected to metabolic labeling with 50 μM 17ODYA or exposed to 0.05% DMSO carrier as control (−17ODYA) for 4 h and lysed. DGKε-Myc was immunoprecipitated with anti-Myc alpaca antibody and subjected to click chemistry reaction with IRDye 800CW-azide. A subset of the 17ODYA-labeled samples were incubated with 1 M HXA for 30 min at 22°C and next diluted twice and incubated for 5 min at 100°C. **(A)** Scheme of the click chemistry procedure. **(B, upper panel**) In-gel fluorescence showing mDGKε-Myc labeling with 17ODYA followed by IRDye-azide, **(B, lower panels)** the efficiency of immunoprecipitation of mDGKε-Myc determined by immunoblotting with mouse anti-Myc antibody. The content of mDGKε-Myc and actin in corresponding input lysates is shown in Fig. 1C of the main text. WT, wild type. Molecular weight markers are shown on the right. **(C)** The extent of mDGKε-Myc palmitoylation. mDGKε-Myc fluorescence was determined by densitometry and normalized against the content of respective mDGKε-Myc variant in immunoprecipitates.

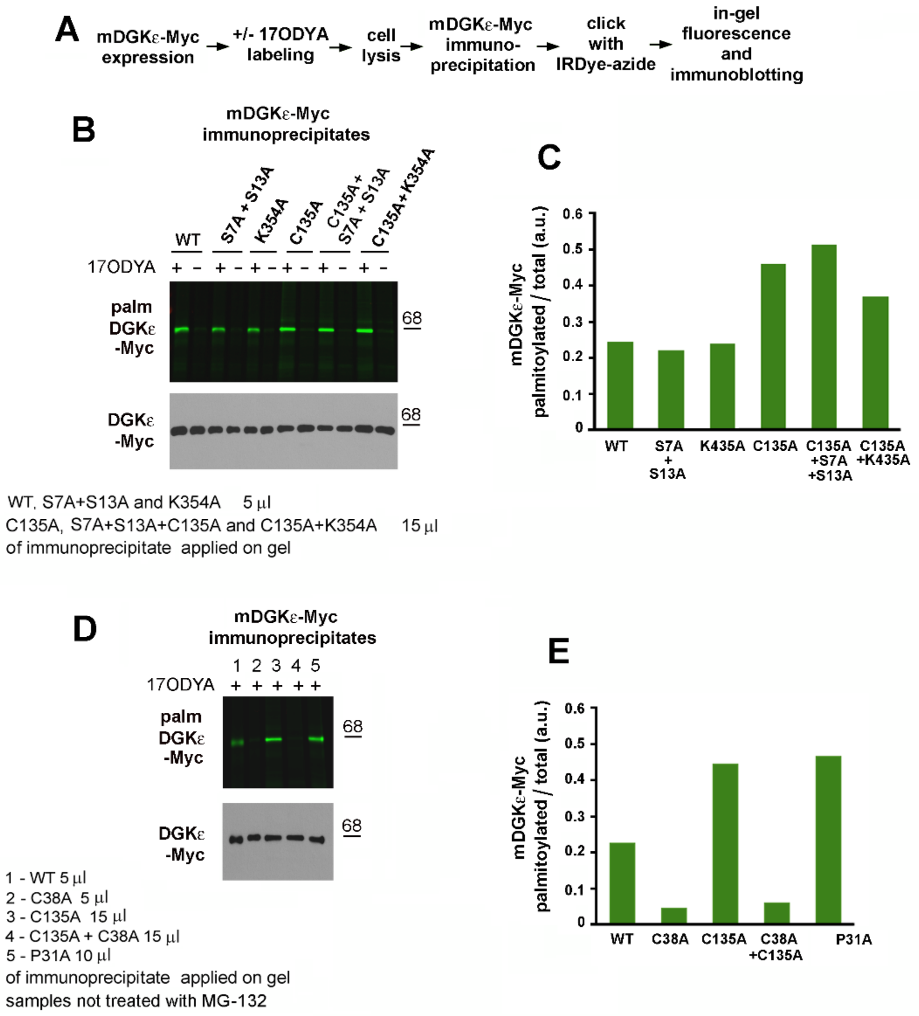


**Supplemental Figure S2. Mouse DGKε is *S*-palmitoylated at Cys38.** HEK293 cells were transfected with plasmid encoding wild type mDGKε-Myc or its indicated mutant forms. After 48 h, cells were subjected to metabolic labeling with 50 µM 17ODYA or exposed to 0.05% DMSO carrier as control (−17ODYA) for 4 h and lysed. mDGKε-Myc was immunoprecipitated with anti-Myc alpaca antibody and subjected to click chemistry reaction with IRDye 800CW-azide. To equalize the content of mDGKe-Myc in samples, the immunoprecipitates were subjected to SDS-PAGE in indicated quantities. **(A)** Scheme of the click chemistry procedure. **(B, D, upper panels)** In-gel fluorescence showing mDGKε-Myc labeling with 17ODYA followed by IRDye-azide, **(B, D, lower panels**) the efficiency of immunoprecipitation of mDGKε-Myc was determined by immunoblotting with mouse anti-Myc antibody. WT, wild type. Molecular weight markers are shown on the right. **(C, E)** The extent of mDGKε-Myc palmitoylation. mDGKε-Myc fluorescence was determined by densitometry and normalized against the content of respective mDGKε-Myc variant in immunoprecipitates.

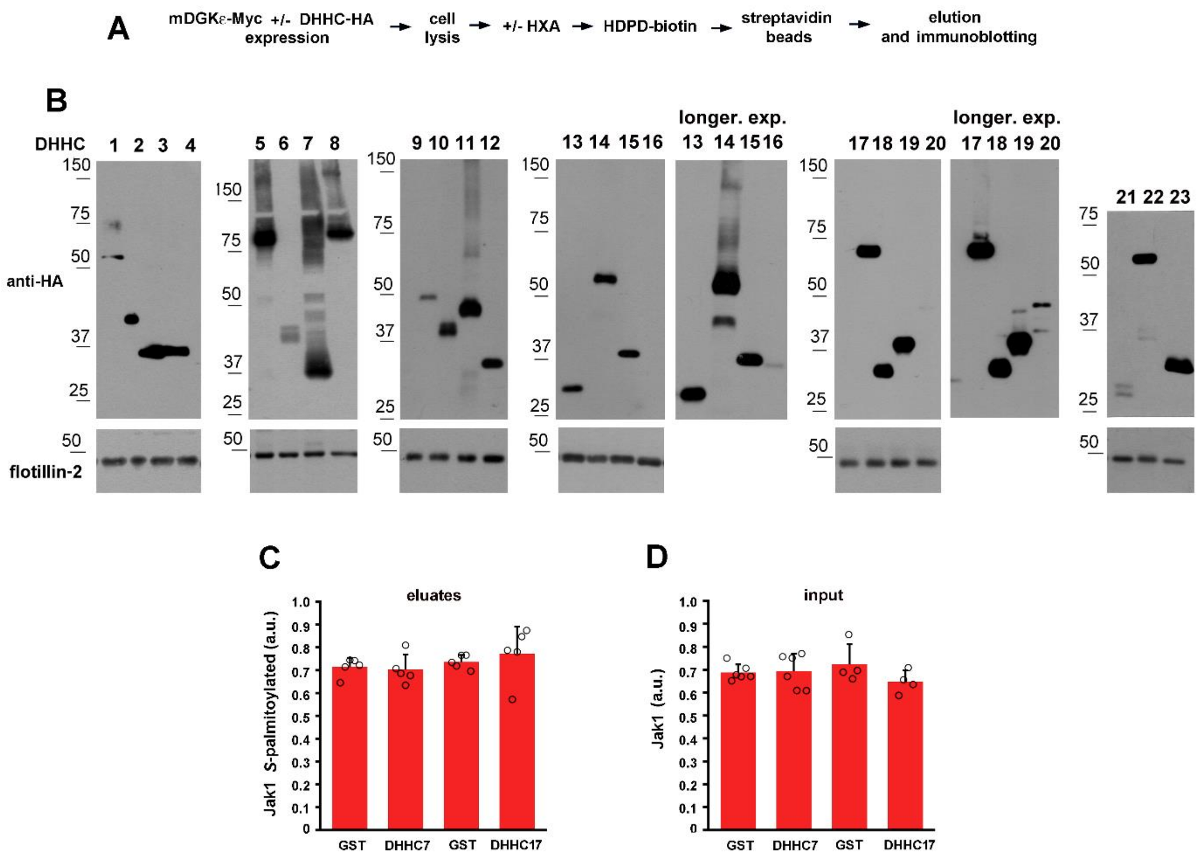


**Supplemental Figure S3. DHHC1-23 are overexpressed in HEK293 cells and do not affect *S*-palmitoylation of endogenous Jak1.** HEK293 cells were co-transfected with plasmid encoding wild type mDGKε-Myc and one of the mouse DHHC1 - 23 *S*-acyltransferases tagged with HA. After 24 h, cells were lysed and proteins were subjected to the ABE procedure involving treatment with hydroxylamine (HXA+) or not (HXA−), biotinylation, and capture of originally *S*-palmitoylated proteins on streptavidin-agarose beads**. (A)** Scheme of the ABE procedure. **(B)** DHHC1-23 expression determined by immunoblotting of input cell lysates with anti-HA-HRP antibody. Endogenous flotillin-2 is shown in lower panels. Shown are results representative of three experiments run in duplicates. Molecular weight markers are on the left. DHHC are numbered after Fukata et al. (25, 34), DHHC10, 11, 13, 22 and 23 correspond to zDHHC11, 23, 24, 13 and 25, respectively. **(C, D)** The level of endogenous *S*-palmitoylated Jak1 **(C)** and the total Jak1 content in input cell lysates **(D)** of cells overexpressing wild type mDGKε-Myc and DHHC7, DHHC17 or GST for control. The content of Jak1 in eluates from streptavidin-agarose beads and in input cell lysates was determined densitometrically. Data are mean ± SD from experiments with DHHC7 and DHHC17 presented in main-text Fig. 3B and used to prepare Figs 3D and 4.

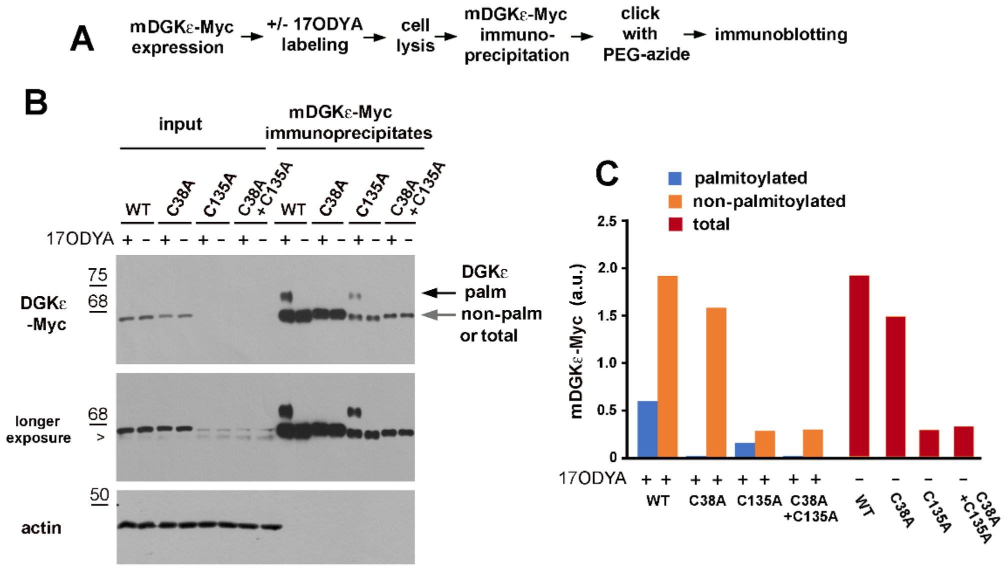


**Supplemental Figure S4. DGKε is *S*-palmitoylated at one cysteine residue.** HEK293 cells were transfected with plasmid encoding wild type mDGKε-Myc or its indicated mutant forms. After 48 h, cells were subjected to metabolic labeling with 50 µM 17ODYA or exposed to 0.05% DMSO carrier as control (−17ODYA) for 4 h and lysed. mDGKε-Myc was immunoprecipitated with anti-Myc alpaca antibody and subjected to click chemistry reaction with PEG-azide. Input lysates and the immunoprecipitates were subjected to SDS-PAGE. **(A)** Scheme of the click chemistry procedure. **(B)** mDGKε-Myc (upper panels) and actin (lower panel) in cell lysates revealed with mouse anti-Myc and mouse anti-actin antibody, respectively. Tagging of 17ODYA-labeled mDGKε-Myc with PEG-azide slows its gel migration and this mobility shift reflects the binding of one PEG-azide to an mDGKε-Myc molecule. Black and gray arrowheads indicate PEGylated (originally palmitoylated) mDGKε-Myc and not modified mDGKε-Myc, respectively. Total mDGKε-Myc content is seen in samples not incubated with 17ODYA that excluded subsequent mDGKε-Myc PEGylation and separation of its labeled (originally palmitoylated) and non-labeled forms. Arrowhead indicates a band recognized unspecifically by the anti-Myc antibody. WT, wild type. Molecular weight markers are shown on the left. **(C)** The extent of mDGKε-Myc palmitoylation. The content of PEGylated (originally palmitoylated), non-palmitoylated and total mDGKε-Myc was determined by densitometry of mDGKε-Myc immunoprecipitates.

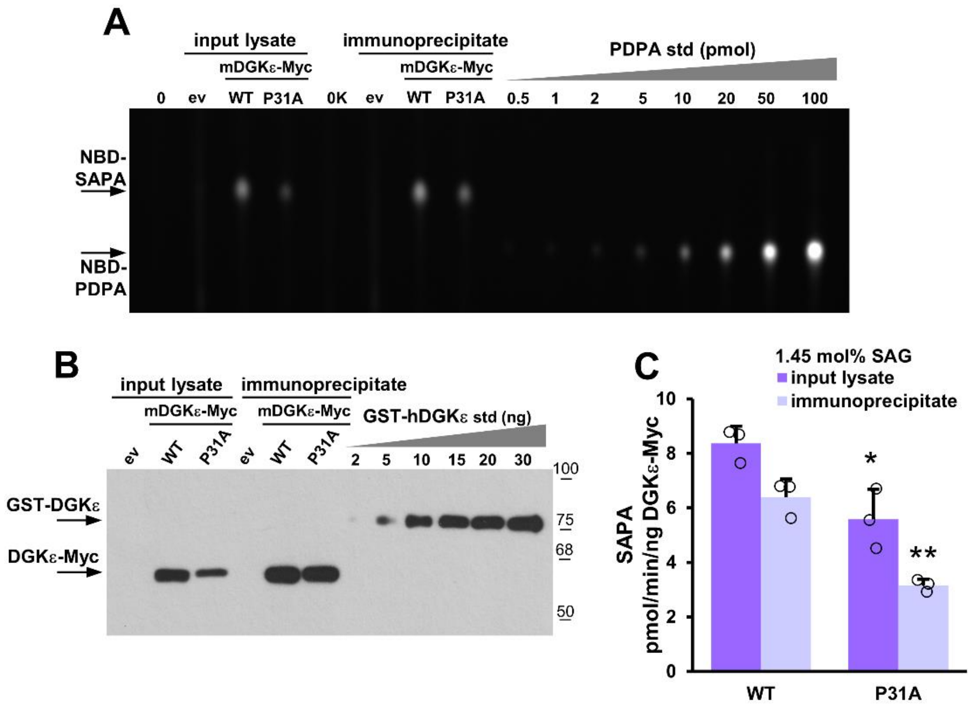


**Supplemental Figure S5. Pro31Ala mutation inhibits DGKε activity.** HEK293 cells were transfected with wild type or Pro31Ala mDGKε-Myc variant or with empty vector (ev) and after 48 h subjected to cell lysis in 1% NP-40 with or without a following mDGKε-Myc immunoprecipitation. The DGKε activity was determined in the cell lysates and mDGKε-Myc immunoprecipitates using a fluorescence assay with mixed micelles of 1.45:2.03 mol% NBD-SAG/SAG:PS. **(A)** Representative TLC separation revealing NBD-SAPA produced. Reactions were carried out using 15 μg of total lysate protein per sample or mDGKε-Myc immunoprecipitates obtained from 75 μg of the lysates. Lipids from 1/25 of the reaction mixture were separated by TLC. NBD-PDPA is used as a standard, it migrates more slowly on TLC than NBD-SAPA. **(B)** Content of indicated overexpressed mDGKε-Myc variants in cell lysates and mDGKε-Myc immunoprecipitates revealed by immunoblotting with sheep anti-DGKε antibody. GST-hDGKε is used as a standard. Three micrograms of total lysate protein and 1/5 of the mDGKε-Myc immunoprecipitate were applied per lane. **(C)** Specific activity of indicated mDGKε-Myc variants calculated after subtraction of the activity of endogenous DGKs determined in control cells (ev). (0) samples devoid of cell lysate, in (0K) supplemented with the Myc-Trap Agarose. WT, wild type. Molecular weight standards are shown on the right. Data are mean ± SD from three experiments. * and **, significantly different at $p < 0.05$ and $p < 0.01$, respectively, from samples of wild type mDGKε-Myc.

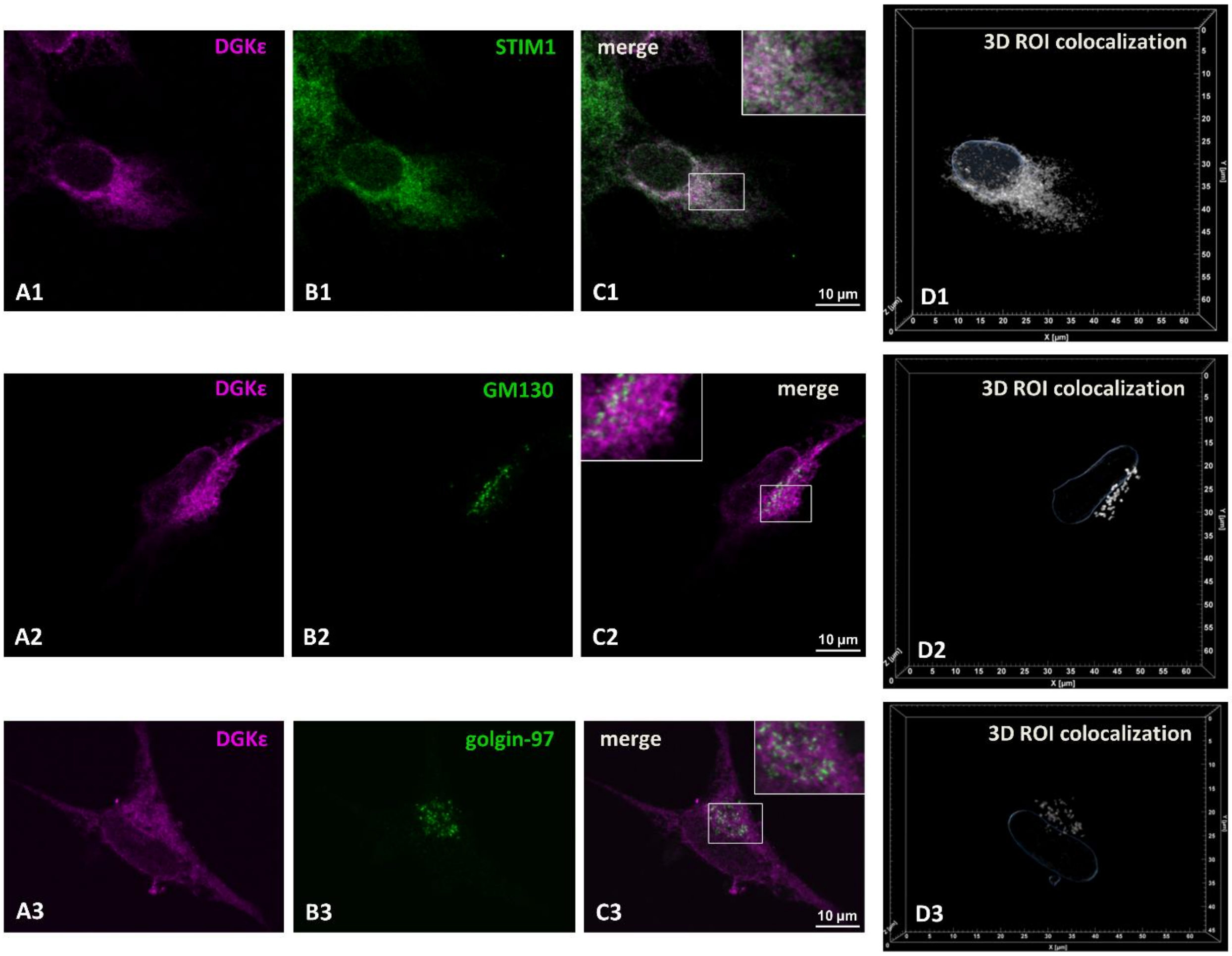


**Supplemental Figure S6. mDGKε-Myc is localized in the endoplasmic reticulum and in the Golgi apparatus.** HEK293 cells were transfected with mDGKε-Myc and after 48 h cells were fixed with 4% paraformaldehyde and permeabilized with 0.05% Triton X-100 (for endoplasmic reticulum staining) or with 0.005% digitonin (for Golgi staining). **(A1-A3)** Localization of mDGKε-Myc, **(B1)** STIM1, **(B2)** GM130, **(B3)** golgin-97. **(C1-C3)** Merged images of mDGKε-Myc and the respective marker protein. Colocalized mDGKε-Myc and marker protein appear white. z-Stack images of ten optical sections taken in the middle of a cell are shown. Insets in **(C1-C3)** show enlarged images of marked fragments. **(D1-D3)** Reconstructed 3D images of two colocalized ROI positive for mDGKε-Myc and STIM1 **(D1)** or GM130 **(D2)** or golgin-97 **(D3)**. Contours of the nucleus detected by Hoechst 33342 staining are shown in blue. ROI as these were used for quantitative analysis of mDGKε-Myc distribution presented in Table 1. mDGKε-Myc (magenta) was visualized with mouse anti-Myc IgG followed by donkey anti-mouse IgG-Alexa647. STIM1, GM130, and golgin-97 (green) were visualized with rabbit anti-STIM1, anti-GM130 or anti-golgin-97 IgG followed by donkey anti-rabbit IgG-FITC. Cells stained according to this protocol are also shown in the main text Fig 7.

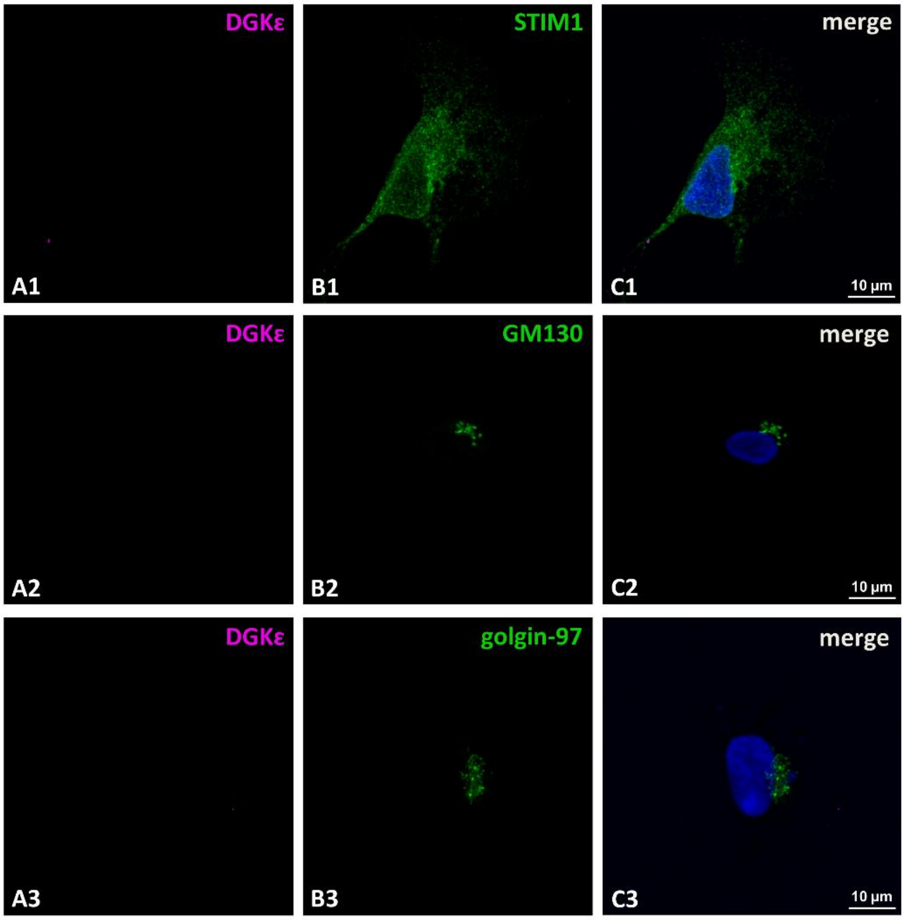


**Supplemental Figure S7. Control experiments indicate specificity of mDGKε-Myc staining.** HEK293 cells were transfected with mDGKε-Myc and after 48 h cells were fixed with 4% paraformaldehyde and permeabilized with 0.05% Triton X-100 (for endoplasmic reticulum staining) or with 0.005% digitonin (for Golgi staining). The procedure of cell staining was like that described in the main text Fig. 7 and supplemental Fig. S6, but without the incubation of cells with anti-Myc antibody. Microscope settings were identical to those used for the colocalization studies. **(A1-A3)** Lack of visible mDGKε-Myc. Localization of **(B1)** STIM1, **(B2)** GM130, **(B3)** golgin-97. STIM1, GM130 and golgin-97 (green) were visualized with rabbit anti-STIM1, anti-GM-130 and anti-golgin-97 IgG followed by donkey anti-rabbit IgG-FITC. All the cells were also incubated with donkey anti-mouse IgG-Alexa647. **(C1-C3)** Merged images with the nucleus seen in blue.

**Supplemental Video 1**

Microscopy images showing co-localization of mDGKε-Myc and STIM1 (image 1), mDGKε-Myc-positive ROI (magenta, image 2), STIM1-positive ROI (green, image 3) and co-localization of mDGKε-Myc- and STIM1-positive ROI (image 4).

**Supplemental Video 2**

Microscopy images showing co-localization of mDGKε-Myc and GM130 (image 1), mDGKε-Myc-positive ROI (magenta, image 2), GM130-positive ROI (green, image 3) and co-localization of mDGKε-Myc- and GM130-positive ROI (image 4).

**Supplemental Video 3**

Microscopy images showing co-localization of mDGKε-Myc and golgin-97 (image 1), mDGKε-Myc-positive ROI (magenta, image 2), golgin-97-positive ROI (green, image 3) and co-localization of mDGKε-Myc- and golgin-97-positive ROI (image 4).
